# *Looplook*: Integrating multiomics refinement and graph clustering for target assignment and functional inference of chromatin regulatory networks

**DOI:** 10.64898/2026.04.03.715516

**Authors:** Ying Zhang, Xingze Huang, Huayu Chen, Long Xie, Ye Chen, Liang Xu

## Abstract

Deciphering target genes regulated by cis-regulatory elements (CREs) is critical for translating genetic and epigenomic findings into clinically actionable insights. However, linking distal CREs to their cognate target genes remains a fundamental challenge due to the limited availability of computational tools for spatial annotation and the oversimplified assignments inherent to conventional topology-only strategies. A flexible framework that integrates 3D proximity with transcriptional output is urgently needed. To address these limitations, we develop *looplook*, an integrated computational framework that bridges 3D chromatin topology with functional genomics to enable accurate, flexible, and user-driven CRE–target gene assignment. *Looplook* provides four core capabilities: (1) robust consensus building for denoising and consolidating replicated or multi-source chromatin loops by employing connected component clustering; (2) bidirectional spatial annotation between 3D chromatin loops and diverse linear genomic features, offering optional graph-based high-order discovery and a linear fallback for gapless network resolution; (3) an expression- or chromatin-aware refinement algorithm that selectively retains functional loops; and (4) automated downstream functional profiling seamlessly integrated with customizable multi-track visualization. Through case studies of the FOSL2 and BRD4 cistromes in liposarcoma cells, we demonstrate that *looplook* outperforms conventional linear annotation methods by integrating chromatin interactions with expression data and chromatin profiles, offering a powerful and valuable framework for distilling experimental omics data into functionally interpretable high-order gene regulation networks. The *looplook* is freely available as an open-source R package, with source code and documentation hosted on GitHub, and will be distributed via the Bioconductor repository.

**Highlights:**

- *Looplook* is an integrative, end-to-end R package for spatial chromatin regulatory network reconstruction.
- *Looplook* supports multiomics integration for expression- and chromatin-aware refinement of target gene annotation, enabling precise linkage of cis-regulatory elements to their cognate target genes.
- *Looplook* distills raw physical chromatin topologies into biologically and/or clinically meaningful gene regulatory networks.

## INTRODUCTION

Precise mapping of cis-regulatory elements (CREs) to their cognate target genes is fundamental to understanding gene expression regulation[1–4]. Dysregulation of these non-coding CREs has been widely implicated in various complex diseases, including cancer and autoimmune disorders, as genome-wide association studies (GWAS) and epigenomic profiling have demonstrated that over 90% of disease-susceptibility variants reside in non-coding genomic regions[5, 6]. Importantly, accurate identification of target genes regulated by distal CREs is essential for translating genetic and epigenomic findings into clinically meaningful insights. A growing body of research, including the exploration of topologically associating domains and oncogenic 3D hubs in human cancers and the comprehensive mapping of disease-risk variants to their target genes[7–10], highlights the immense potential of these spatial regulatory architectures in empowering therapeutic target prioritization and precision medicine applications[11].

Establishing robust CRE–target gene linkages remains a long-standing goal in the field of experimental biology[2–4]. Advances in chromosome conformation capture technologies, such as high-throughput/resolution chromosome conformation capture (Hi-C), in situ Hi-C followed by chromatin immunoprecipitation (HiChIP), chromatin interaction analysis by paired-end tag sequencing (ChIA-PET), and proximity ligation-assisted chromatin immunoprecipitation sequencing (PLAC-seq), have provided critical insights into spatial chromatin architecture and substantially improved the accuracy of CRE–target gene assignment[12–18]. Clustered regularly interspaced short palindromic repeats (CRISPR)-based perturbations further offer direct functional evidence, but they are inherently low-throughput and heavily reliant on pre-identified chromatin interactions[19, 20].

Complementing experimental efforts, a growing repertoire of computational tools has been developed to annotate CRE–target linkages. Conventional annotation methods (e.g., *ChIPseeker*, *rGREAT*) primarily rely on linear genomic proximity, assigning CREs to the nearest genes or those within a fixed scanning window[21, 22]. While conceptually simple and computationally efficient, these approaches often yield high false-negative rates by failing to capture distal interactions mediated by 3D chromatin looping. Indeed, distal enhancers frequently bypass linearly adjacent genes via chromatin loops to directly regulate targets megabases away[11]. High-resolution 3D chromatin data are rapidly expanding, but dedicated 3D-guided annotation frameworks has lagged behind, with only a limited number of tools currently incorporating spatial chromatin conformation information to bridge this gap. For instance, the *Activity–by–Contact* (*ABC*) model utilizes built-in databases combining chromatin accessibility, Hi-C contacts, and epigenetic signals to score enhancer–gene pairs[19]. The 3D chromatin module in *FUMA* is tailored primarily for annotating GWAS variants[23]. Furthermore, while most 3D loop callers (e.g., *FitHiChIP*, *hichipper*, *PSYCHIC*, *cLoops2*, *MUSTACHE*, *HiCExplorer*, *DLoopCaller*, and *UnionLoops*) excel at delineating spatial chromatin contacts, their primary focus typically does not extend to comprehensive downstream functional annotation[24–32]. Consequently, downstream analysis frequently involves a combination of custom scripts, command-line suites (e.g., *bedtools*), and foundational Bioconductor parsers (e.g., *GenomicInteractions*) to facilitate basic geometric intersections for regulatory assignment[33, 34]. Reliance solely on coordinate overlaps often yields elevated false-positive rates; moreover, the fragmented and non-standardized workflows typically employed can limit the reliability and stability of the resulting biological mechanistic interpretations.

Alternatively, several studies implemented regression or machine learning approaches (e.g., *FOCS*, *TargetFinder*, *SCEG-HiC*, *SPEID*, *EP2vec*, *DeepPHiC*, and *ENCODE-rE2G*) to infer/predict enhancer–promoter (E–P) links[2, 35–40]. Although these existing tools incorporating 3D chromatin information have significantly improved target prediction accuracy, they still face inherent limitations across data compatibility, algorithmic design, and analytical completeness. First, regarding data input, many tools are restricted to rigid, precompiled data ecosystems. Because CRE function is highly context-dependent across species, lineages, and cell states, static databases lack the flexibility to incorporate user-defined, context-matched multiomics data. Second, from an algorithmic perspective, current strategies oversimplify the biological complexity of gene regulation. On one hand, they predominantly rely on static physical co-localization, which does not guarantee genuine transcriptional activation; without integrating dynamic transcriptome profiles, these approaches frequently generate false-positive assignments[41]. On the other hand, by strictly focusing on isolated pairwise contacts, traditional methods fail to resolve higher-order topological architectures. Biologically, gene regulation can operate through modular and “multi-hop” spatial interactions, where topological connectivity (hubness) is intrinsically coupled with robust transcriptional outputs[7, 42]. Therefore, incorporating these multi-hop dependencies is increasingly recognized as essential for a more holistic understanding of spatial regulatory networks. Third, concerning analytical workflows, most tools output target gene lists without integrated downstream functional analyses, such as transcription factor (TF) motif scanning, Protein–Protein Interaction (PPI) network construction, pathway enrichment, and customizable multi-track visualization. Together, these bottlenecks highlight a critical unmet need for a versatile and integrated computational framework.

To systematically address these limitations, we developed *looplook*, a flexible, end-to-end computational suite designed to consolidate heterogeneous 3D chromatin topologies, resolve higher-order regulatory connectivity, and deliver integrated functional annotation. The core advantage of *looplook* is driven by three key features: (i) expression-aware topological reclassification, which suppresses transcriptionally silent false positives by re-annotating locally silent promoters and gene bodies as enhancer-like elements (named as eP and eG, respectively) while preserving graph connectivity; (ii) chromatin-aware anchor refinement, which leverages orthogonal histone modification and chromatin accessibility data to resolve ambiguous eP/eG classifications into definitive anchor types, including dual-function promoter/enhancer elements; and (iii) robust topological reconstruction through connected component clustering and multi-hop network diffusion. By circumventing the constraints of closed data ecosystems, our framework empowers researchers to transition from simplistic static target assignments to dynamic, high-confidence inference and curation of functional regulatory network.

## RESULTS

### Implementation and key features of *looplook*

*Looplook* provides an end-to-end analytical pipeline driven by a core computational architecture comprising five interconnected modules (**Figure 1**): (1) Replicate Consolidation & Multi-Source Consensus; (2) 3D-Guided Annotation & Spatial Bridge Mapping; (3) Expression- and Chromatin-Aware Refinement; (4) Functional Profiling; and (5) Visualization. Key features of *looplook* are summarized as below.

**Figure 1.**
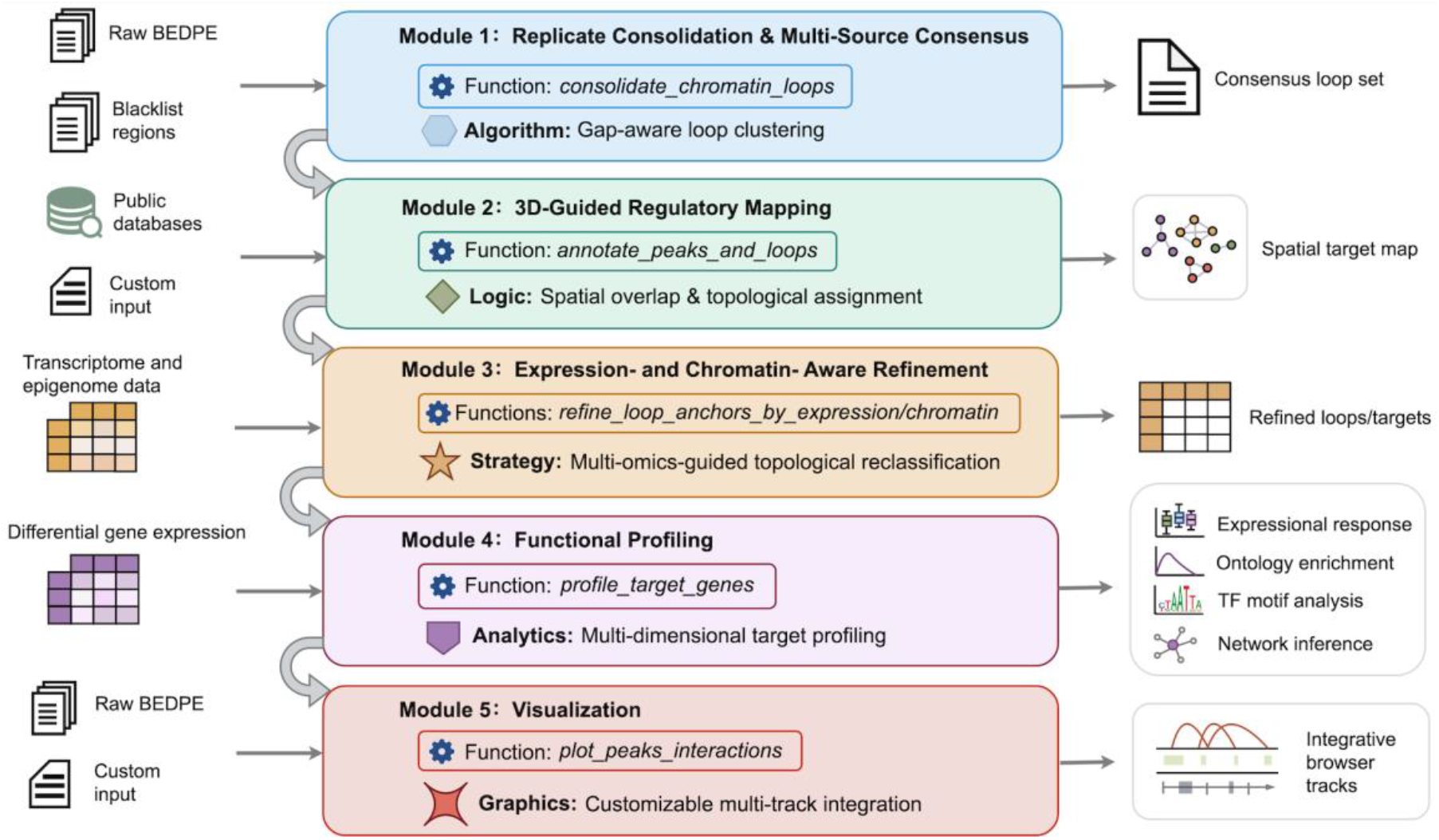
Overview of the *looplook* analytical framework and workflow. The pipeline consists of five integrated modules that process multi-omics data: (1) Replicate Consolidation & Multi-Source Consensus, which integrates raw spatial chromatin loops from multiple sources or replicates; (2) 3D-Guided Annotation & Spatial Bridge Mapping, which links user-defined genomic features to cognate target genes; (3) Expression- and Chromatin-Aware Refinement, which filters transcriptionally silent interactions based on the available expression data; (4) Functional Profiling, which performs multi-dimensional characterization of target genes and their associated pathways; and (5) Visualization, which enables integrated exploration of chromatin topologies and multi-track visualization of epigenetic data.

### Feature 1: 3D topology consolidation and graph instantiation

For 3D chromatin conformation data derived from multiple biological replicates (typically in BEDPE format), the *consolidate_chromatin_loops* function in Module 1 applies a topology-merging engine based on connected component clustering (**Figure S1**). To mitigate batch effects and technical noise, this engine supports three customizable consolidation modes: intersect mode (retains loops with fully overlapping anchors across all replicates), consensus mode (default, retains loops supported by the majority of replicates), and union mode (retains all detected loops).

### Feature 2: 3D-guided annotation and spatial bridge mapping

After overall graph instantiation, the *annotate_peaks_and_loops* function in Module 2 processes user-input tabular omics datasets in parallel, including transcriptomics, chromatin accessibility, single nucleotide polymorphisms (SNPs) or genetic variants annotated by GWAS, and protein–DNA interaction information derived from chromatin immunoprecipitation sequencing (ChIP-seq), cleavage under targets and tagmentation (CUT&Tag), or cleavage under targets and release using nuclease (CUT&RUN) (**Figure 2A,B**). At this stage, the system abstracts the underlying spatial topology into an undirected spatial graph model:

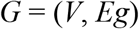

**Figure 2.**
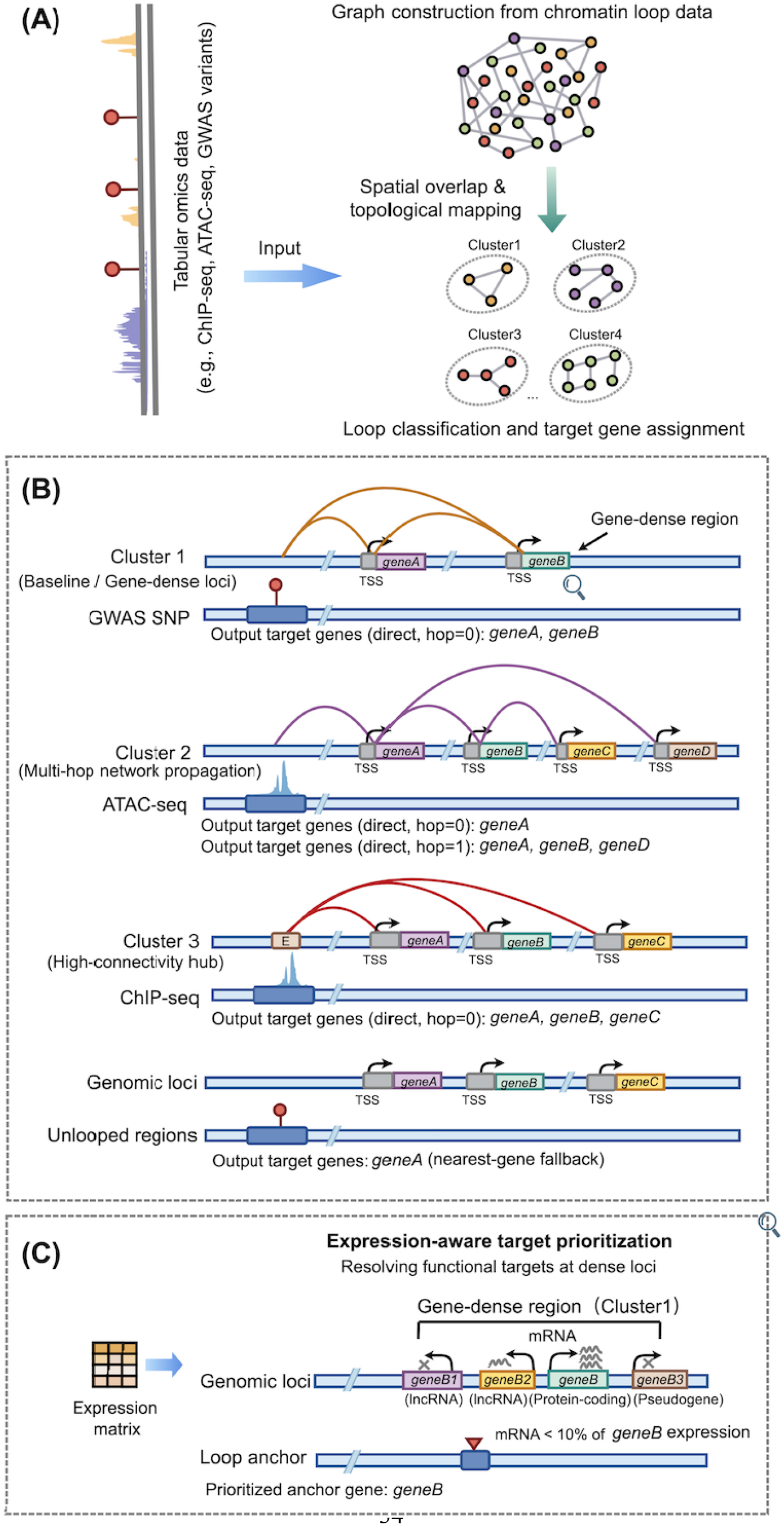
Graph-theoretic modeling and multi-dimensional target annotation strategies in *looplook*. (A) Integration of user-input tabular omics data and 3D chromatin loops into an undirected spatial graph, partitioned into independent topological clusters. (B) Schematic representations of diverse annotation schemes: direct spatial mapping (Cluster 1); higher-order multi-hop network diffusion bridging indirect targets (Cluster 2, controlled by the *neighbor_hop* parameter); hub identification for high-connectivity regulatory centers (Cluster 3); and a linear fallback mechanism ensuring coverage for regions lacking loops (Cluster 4). (C) Schematic of the expression-guided refinement module which helps to resolve mapping ambiguities at gene-dense regions by filtering out transcriptionally silent or lower-priority candidates.

Here, the vertex set *V* represents all chromatin loop anchors (nodes) in the input interaction network. Anchors overlapping with input regulatory or variant signals are classified into the regulatory node subset *Vr*. Those overlapping with gene promoters are assigned to the promoter node subset *Vp*. These subsets are not necessarily mutually exclusive, as an anchor may simultaneously overlap a regulatory feature and a promoter. Anchors devoid of these features are retained as unannotated topological nodes. The edge set *Eg* denotes direct physical contacts mediated by chromatin loops (edges). Following node annotation, edges in *Eg* are dynamically mapped and categorized into functional topological interactions according to the genomic identities of the connected loci, including E–P, promoter–promoter (P–P), and enhancer–gene body (E–G) interactions.

For a regulatory node (*v*∈*V_r_*), its *k*-hop spatial neighborhood is further defined as

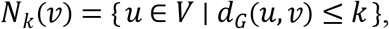

where *d_G_*(*u*,*v*) denotes the shortest-path distance between nodes *u* and *v* in graph *G*. This neighborhood definition provides the formal basis for topology-aware target expansion, allowing regulatory signals to be propagated from directly overlapping anchors to spatially connected neighboring nodes within a user-defined hop distance.

### Feature 3: Context-aware refinement

#### 1. Expression-aware topological reclassification (E-refined mode)

To resolve mapping ambiguities arising from chromatin loop anchors that span gene-dense regions, Module 3 replaces traditional static filtering with a topological reclassification strategy guided by integrated expression information (e.g., matched transcriptomic data) (**Figure 2C**). This strategy demonstrates robust denoising and reconstruction capabilities when processing baseline and gene-dense loci (as exemplified in Cluster 1). It executes a three-step cascading mechanism: (i) integrating a transcript abundance matrix to pre-filter transcriptionally silent genes; (ii) prioritizing candidate targets by user-defined gene biotype (e.g., protein-coding > non-coding); and (iii) applying a dominant expression tiebreaker. For instance, when confronted with highly overlapping target clusters, the algorithm evaluates the local transcriptional neighborhood and systematically prunes baseline noise by retaining only those candidate genes whose expression levels exceed 10% of the maximum transcript abundance within that anchor.

Beyond prioritizing active targets and resolving local assignment conflicts, Module 3 further addresses the regulatory potential of transcriptionally silent genomic anchors (e.g., promoters and gene bodies) through dynamic topological reclassification. In light of accumulating evidence that both gene promoters and internal gene bodies can exert enhancer-like activity[3, 43–45], the algorithm accounts for the possibility that these elements may function as distal regulators despite the absence of local transcription. To operationalize this, the *refine_loop_anchors_by_expression* function does not merely sever network connectivity for genomic anchors that are transcriptionally silent (expression below a defined threshold). Instead, it deploys a dynamic topological reclassification, through which silent promoter and gene bodies are re-annotated as putative distal enhancer-like elements (hereafter referred to as P-to-eP and G-to-eG reclassification, respectively) capable of regulating other target genes. Computationally, this strategy models the biological context where local transcriptional activity is not detected while the 3D physical scaffold persists to support the expression of distal target genes. Following this attribute shift, the original edge set dynamically evolves, reconfiguring traditional functional edges (e.g., E–P, P–P, and E–G) into enhancer-like topological connections (e.g., E–eP, eP–P, and eG–P) (**Figure 3**).

**Figure 3.**
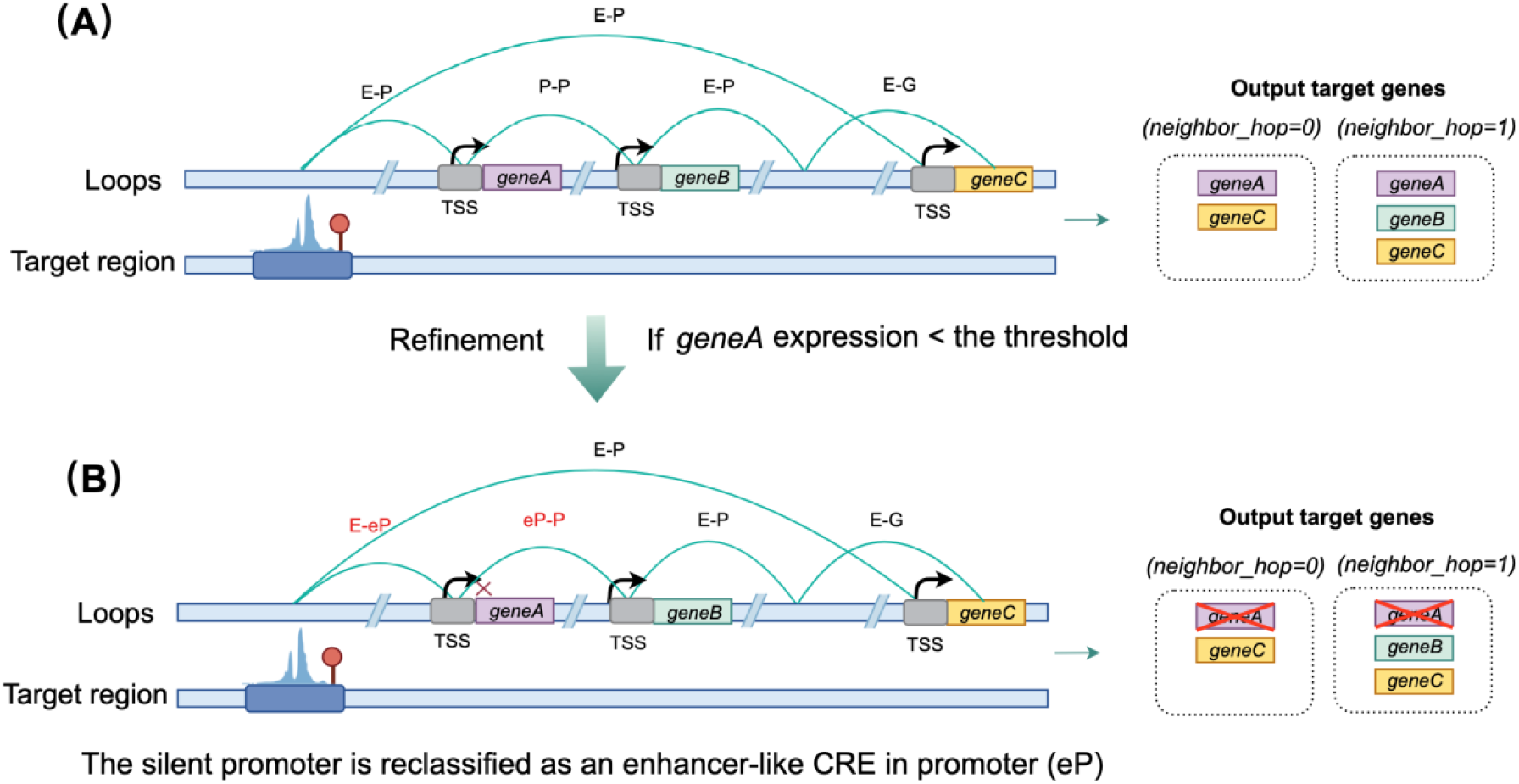
Schematic of expression-guided topological reclassification. (A) In a typical topology-based annotation, regulatory signals (e.g., from a distal enhancer) are mapped to all physically connected promoters (e.g., *geneA*, *geneB*, *geneC*). (B) Upon integrating transcriptomic data, *looplook* identifies *geneA* as transcriptionally silent (expression < threshold). Rather than completely removing the promoter node, which would sever indirect connections, the reclassification algorithm alters its attribute to an enhancer-like element (eP) and subsequently reconfigures E–P and P–P edges into E–eP and eP–P edges (highlighted in red), respectively. The output target list is refined (excluding the silent *geneA*) while preserving the higher-order spatial scaffold for multi-hop signal transmission (e.g., successfully retaining *geneB* at *neighbor_hop* = 1).

#### 2. Chromatin-aware anchor refinement (C-refined mode)

The *refine_loop_anchors_by_chromatin* function provides orthogonal validation and reclassification of loop anchor types using direct histone modification and chromatin accessibility evidence. This module addresses a fundamental limitation of both positional annotation and expression-based refinement: neither can determine the chromatin identity of a given locus. A promoter annotated as silent by expression filtering (eP) may in fact harbor active enhancer marks, while a distal element classified as E may carry promoter-associated signals. Chromatin-aware refinement resolves these ambiguities by integrating additional chromatin profiles: H3K4me1 (enhancer-associated), H3K4me3 (promoter-associated), H3K27ac (active enhancer/promoter), ATAC-seq (chromatin accessibility), and H3K27me3 (Polycomb-mediated repression).

For chromatin state inference, each candidate anchor is first evaluated for peak overlap using cascade-filtered peak-to-anchor matching with configurable gap tolerance (*anchor_gap*, default 200 bp), minimum physical overlap (*anchor_min_overlap*, default 100 bp), and minimum overlap fraction (*anchor_min_frac*). A chromatin state is then assigned to each anchor based on its mark combination, with priority given to the most informative state (**Table S1**).

Anchor-type reclassification applies a deterministic rule set that maps each combination of positional type and chromatin state to an updated classification. The anchor-type reclassification rules, evaluated in decreasing order of priority, are summarized in **Table S2-3**. The minimal required inputs are H3K4me1 and H3K4me3; H3K27ac, ATAC, and H3K27me3 are optional but can improve confidence.

Following expression-driven refinement that clears the gene symbol of a silent anchor (P-to-eP or G-to-eG conversion), chromatin evidence may subsequently re-assign the anchor back to P, G, or “Dual”. Under these circumstances, the original gene symbol must be restored so that the reclassified anchor contributes to promoter-centric statistics and target gene assignment. The chromatin-aware refinement module recovers original gene symbols from the internal anchor state attributes preserved during the initial annotation step. Additionally, when “*recompute_targets* = TRUE” (default), the module rebuilds the peak-to-gene linkage table using updated anchor types, producing chromatin-aware *target_annotation* and *target_gene_links* outputs with full provenance tracking. Each entry in *target_gene_links* is annotated with columns: *anchor_type_before_chromatin*, *anchor_type_after_chromatin*, *chromatin_confidence*, and c*hromatin_evidence*, enabling row-level auditing of chromatin-guided reclassification events.

#### 3. Integration of expression- and chromatin-aware refinement (I-refined mode)

The expression-aware and chromatin-aware refinement modules address orthogonal biological questions: transcriptional activity versus chromatin identity. When both RNA-seq and ChIP-seq datasets are available, the refinement procedure follows a defined sequential workflow in which expression-aware refinement is performed first, followed by chromatin-aware refinement. Expression refinement first evaluates transcriptional support for candidate anchor–gene assignments and marks transcriptionally unsupported or ambiguous anchor states as tentative hypotheses (eP/eG), followed by chromatin refinement that validates or revises these hypotheses into definitive anchor types (E, P, G, or Dual). Columns derived from expression-aware refinement process (*Active_Target_Genes*, *Retained_In_Functional_Network*, *Refinement_Action*) remain unchanged throughout chromatin refinement, ensuring that downstream functional profiling operates on the expression-validated gene set while benefiting from chromatin status-assisted anchor classifications.

### Feature 4: Multi-hop network diffusion and adaptive hubs

The aforementioned P-to-eP reclassification preserves graph connectivity, laying the foundation for higher-order multi-hop network diffusion (exemplified by Cluster 2 in **Figure 2**). Users can transcend direct physical contact constraints by tuning the *neighbor_hop* parameter. For instance, setting “*neighbor_hop* = 1” allows regulatory signals to propagate beyond the primary target (e.g., *geneA*) and to reach indirectly connected loci (e.g., *geneB*, *geneC*, and *geneD*) via the network topology. Crucially, even when an intermediate node undergoes P-to-eP reclassification (e.g., a transcriptionally silent *geneA*), the regulatory signal can still be relayed along this eP-mediated scaffold to secondary nodes deeper within the network (illustrated by retention of *geneB* at “*neighbor_hop* = 1” in **Figure 3**). Concurrently, to evaluate the topological significance of distal elements, *looplook* adaptively detects high-connectivity hubs (**Figure 2**, Cluster 3). If the degree centrality (i.e., loop connectivity count) of a regulatory anchor exceeds a user-specified genome-wide distribution threshold (controlled by the *hub_percentile* parameter), the node is formally designated as a key regulatory hub orchestrating multiple downstream promoters.

### Feature 5: Smart fallback for gapless annotation

To handle genomic features such as distal variants or orphan peaks lacking 3D chromatin loop support, *looplook* implements a “Smart Fallback” procedure (**Figure 2**, Cluster 4). The algorithm automatically switches to a linear proximity-based search and assigns each input feature to its nearest transcriptionally active gene. This scheme achieves comprehensive annotation coverage for all input signals and prevents loss of biologically relevant information within the multiomics integration workflow. The fully annotated and reclassified network is subsequently passed to the downstream functional profiling and visualization modules for automated multi-dimensional exploration.

### Feature 6: Functional profiling and multi-track visualization

As an end-to-end analytical framework, *looplook* encapsulates fully automated downstream functional profiling and comprehensive visualization within one unified workflow. In Module 4, the *profile_target_genes* function executes multi-dimensional target decoding, seamlessly linking annotated gene sets to transcriptional dynamics analysis, target signature enrichment, and core regulatory network inference pipelines. Module 5 provides high-quality visualization through the *plot_peaks_interactions* function, built upon a customizable multi-track integration architecture. This module supports vertical alignment and rendering of user-supplied epigenetic or variant signals, 3D chromatin loop topologies, and linear gene transcript models within a shared genomic coordinate system. The resulting integrative browser-style tracks intuitively reconstruct spatial regulatory landscapes at complex loci, substantially bridging abstract graph-theoretic networks to concrete mechanistic biological hypotheses.

### Methods comparison

*Looplook* introduces an elegantly architected framework for CRE-to-target gene assignment. Using graph-based clustering, the algorithm standardizes, denoises, and merges multi-source 3D chromatin loops to construct high-confidence regulatory networks. To rigorously reduce false positives and improve target prediction accuracy, the framework implements expression-guided refinement alongside dynamic topological reclassification. Furthermore, it identifies distal regulatory hubs via multi-hop network diffusion and achieves exhaustive annotation coverage via the smart linear fallback mechanism. Moving beyond topological mapping, *looplook* integrates TF motif scanning, PPI network construction, pathway enrichment analysis, and publication-grade multi-track visualization within a unified analytical environment. A side-by-side comparison of core capabilities between *looplook* and current state-of-the-art annotation tools is summarized in **Table 1**.

**Table 1.** Comparison of core features between *looplook* and current state-of-the-art genomic annotation tools.

| Feature/Capability | *ChIPseeker* | *GREAT* | *Genomic-Interactions* | *FUMA* | *ABC Model* | *looplook* |
| --- | --- | --- | --- | --- | --- | --- |
| Primary annotation scheme | 1D | 1D | 3D | 3D | 3D | 1D & 3D |
| User-input 3D interactome | - | - | √ | - | √ | √ |
| User-input omics data | √ | √ | - | - | √ | √ |
| Graph-based topological consensus | - | - | - | - | - | √ |
| Expression-/Chromatin-guided refinement | - | - | - | P* | - | √ |
| Topological reclassification (eP/eG) | - | - | - | - | - | √ |
| Multi-hop network diffusion | - | - | - | - | - | √ |
| Smart linear fallback mechanism | - | - | - | - | - | √ |
| Functional inference | - | √ | - | - | - | √ |
| Multi-track integrative visualization | - | - | - | - | - | √ |
| Local execution (data privacy) | √ | - | √ | - | √ | √ |
| Ecosystem | R | Web | R | Web | Python | R |

Note: “1D” refers to linear proximity-based assignment; “3D” refers to assignment guided by spatial chromatin topology. “√” denotes full native support; “-” denotes not supported; “P\*” denotes partial support (*FUMA* relies on precompiled reference datasets rather than user-supplied data).

### Applications and examples

Seminal studies have established that dedifferentiated liposarcoma (DDLPS) cells are critically dependent on both bromodomain and extraterminal (BET) proteins and FOSL2, with BRD2/3/4 maintaining the transcriptional output of active enhancer networks and the core transcriptional regulatory circuitry comprising FOSL2, MYC, and RUNX proteins[46–49]. We showed that BET-targeting proteolysis-targeting chimeras, ARV-825 and a structurally different compound ZBC-260, effectively depleted intracellular levels of BRD2/3/4 and FOSL2 proteins (**Figure S2**). This experimentally validated gene regulatory program serves as an ideal system for evaluating *looplook*’s utility and performance. We therefore applied *looplook* to a multiomics dataset from a DDLPS cell line LPS141 (**Table S4**), comprising newly generated and published RNA-seq for transcriptional responses to BRD4 degradation, ChIP-seq for FOSL2 and BRD4, histone mark ChIP-seq and ATAC-seq for chromatin status, and H3K27ac HiChIP for 3D chromatin interactions[47, 50]. *Looplook* efficiently (1) reconstructs 3D transcriptional regulatory networks, (2) improves enhancer–target gene linkage, and (3) outperforms conventional linear assignment in capturing cistrome dynamics under pharmacological perturbation.

### Case study 1: *Looplook* uncovers the functional cistrome of BRD4 in LPS141 cells

Genome-wide H3K27ac HiChIP revealed loop anchors distributed across all autosomes (**Figure S3A**). Further leveraging expression data to refine target annotation (E-refined mode), *looplook* revealed extensive P–P (33.0%) and gene body–promoter (G– P, 22.5%) interactions as the dominant topological configurations (**Figure S3B**). In line with the observation that over 40% of H3K27ac HiChIP loop anchors resided in non-promoter elements, including intronic (18.4%) and distal intergenic regions (8.5%) (**Figure S3C**), *looplook* demonstrated that 78.2% of total BRD4 binding events (*n* = 14,033) were physically tethered within E–P loops, and that 86.5% of them were associated with active transcription (**Figure 4A**). Pharmacological depletion of BRD2/3/4 proteins with ZBC-260 elicited strong perturbation of cellular transcriptome (**Figure S3D**). Furthermore, integrative analysis of BRD4 ChIP-seq and H3K27ac-associated 3D looping/connectivity, *looplook* refined linear annotations into functional regulatory states, converting distal intergenic and intronic peaks into enhancers (E, 48.2%) or dual-identity promoter-/enhancer-like elements (Dual, 8.5%) (**Figure S3E**). Chromatin profiling confirmed that reclassified anchors engaging enhancers were enriched for active marks, including H3K4me1 (88%–100%), H3K27ac (100%), and Tn5 accessibility (82%–98%), whereas promoter-associated anchors maintained robust H3K4me3 occupancy (100%) without repressive H3K27me3 (**Figure S3F**).

**Figure 4.**
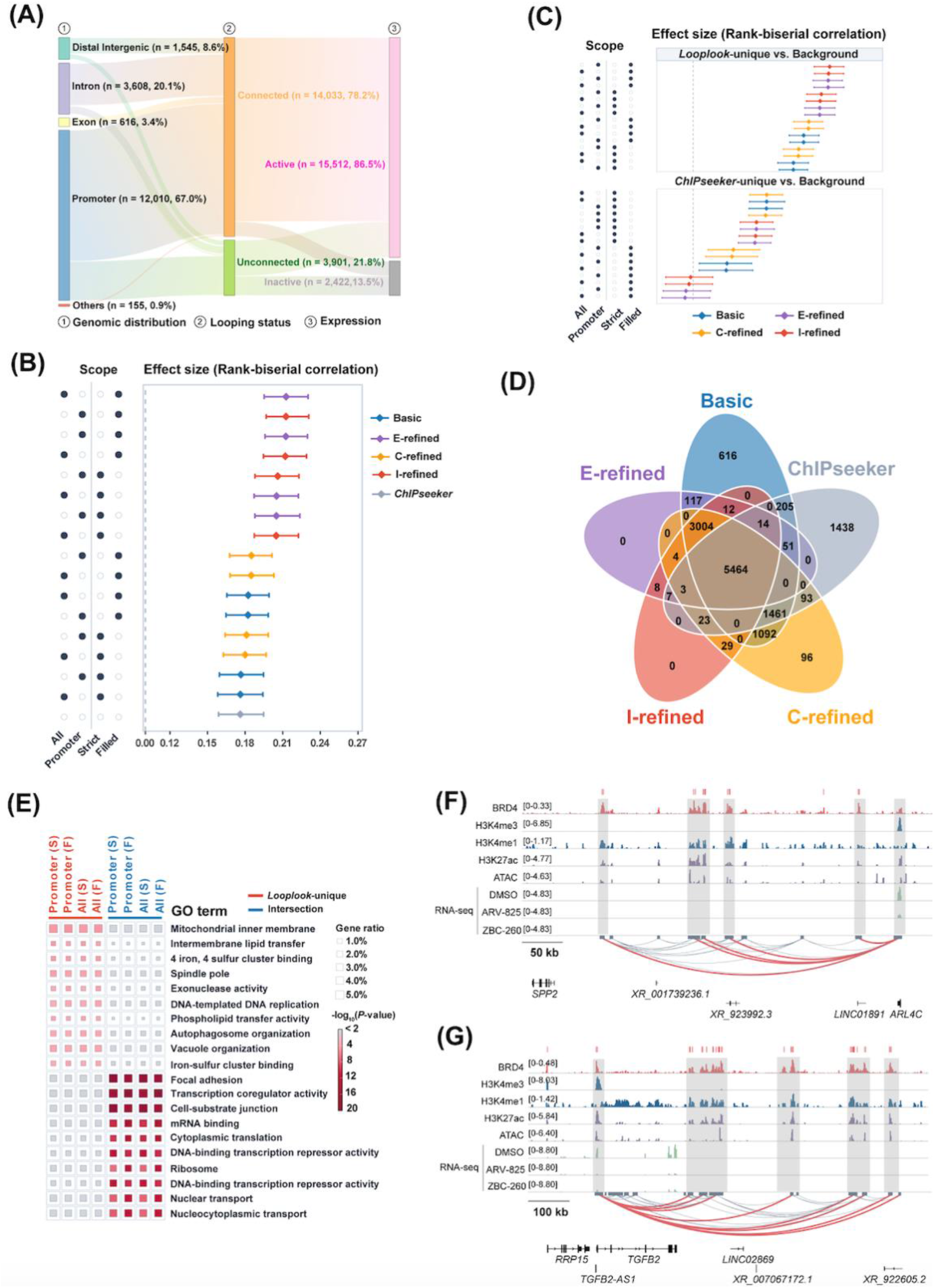
Comparative benchmarking of *looplook* against the conventional linear annotation approach in decoding the BRD4 cistrome. (A) Stacked bar plots showing the genomic distribution of BRD4 peaks, looping status (Connected vs. Unconnected), and target gene transcriptional activity (Active vs. Inactive). (B) Effect size estimates (rank-biserial correlation) for BRD4 target gene sets identified by *looplook* across four refinement configurations (Basic, E-refined, C-refined, and I-refined) and four peak−target annotation scopes (All: all peaks; Promoter: promoter-associated peaks; Strict: loop-engaged targets, Filled: loop-engaged targets plus fallback-filled targets), compared with the conventional linear nearest-TSS method (*ChIPseeker*) under ZBC-260 treatment. Data points represent effect size estimates with error bars indicating 95% confidence intervals. (C) Comparative effect size distributions across selected target gene subsets (*looplook*-unique vs. Background; *ChIPseeker*-unique vs. Background) across indicated annotation modes. (D) Venn diagram showing the overlap and unique BRD4 target genes identified by *looplook* (Basic, E-refined, C-refined, and I-refined modes) versus *ChIPseeker*. (E) GO enrichment analysis comparing *looplook*-unique, commonly annotated (intersection), and *ChIPseeker*-unique BRD4 target genes. Dot sizes represent gene ratios; colors indicate statistical significance (−log_10_ *P*-value) across multiple peak−target scopes (S, strict; F, fallback-filled). (F,G) Linkage of distal BRD4-bound regulatory elements to the *ARL4C* and *TGFB2* loci in LPS141 cells.

Next, we subjected *looplook* to comparative benchmarking against the conventional nearest transcription start site (TSS) annotation strategy (*ChIPseeker*), employing four refinement configurations (Basic, E-refined, C-refined, and I-refined) for looped targets, optionally supplemented with Fallback annotation for unlooped BRD4 peaks (**Figure 4B–D**, and **Table S5**). Effect size evaluations demonstrated that *looplook*-assigned BRD4 target genes under E-/C-/I-refined modes generally exhibited stronger ZBC-260 responsiveness than *ChIPseeker*, with or without Fallback annotation (**Figure 4B**), suggesting that *looplook* improves biological relevance. Notably, expression-aware refinement made a substantial contribution to *looplook*’s performance in capturing ZBC-260-responsive BRD4 targets, compared with chromatin-aware refinement (**Figure 4C, Figure S4A**). Importantly, this performance advantage was stable and independently reproducible upon ARV-825-mediated BET degradation (**Figure S4B,C,** and **Table S5**). Iterative Gene Set Enrichment Analysis (GSEA) permutations and cumulative moving average trajectories of normalized enrichment scores (NES) revealed that *looplook*-unique gene sets exhibited markedly smaller and sustained negative NES values than *ChIPseeker*-unique counterparts across both ZBC-260 and ARV-825 conditions (**Figure S4D–G**), demonstrating the superior robustness of *looplook* over *ChIPseeker* in capturing biologically relevant targets.

Notably, *looplook* and *ChIPseeker* commonly annotated over 5,500 BRD4 targets, while the refinement strategies (E-refined, C-refined, and I-refined) exhibited distinct impacts on the identification of new BRD4 targets by *looplook* (**Figure 4D**). Analysis of the distance between BRD4 peaks and their assigned target TSS revealed that *looplook*-unique targets engaged long-range interactions, with intervals ranging from 10 kb to over 100 kb (**Figure S5A,B**). Specifically, *looplook* enabled the annotation of an expanded set of BRD4 target genes associated with BRD4 occupancy at distal CREs within enhancers, enhancer-like promoters, or gene bodies, and likely regulated through chromatin interactions (**Figure S5C–E**). By contrast, *ChIPseeker*-unique annotations were intrinsically constrained to proximal promoters. These results highlight the competence of *looplook* in identifying distal BRD4 targets.

In addition to target annotation, *looplook* integrates several streamlined bioinformatics tools, including motif analysis, pathway enrichment analysis, PPI network analysis, and a genomics viewer. The motif analysis function enables exploration of TF binding site enrichment across selected peak or anchor sets. For instance, d*e novo* motif analysis of loop anchor-associated BRD4 peaks indicated significant enrichment of DNA binding motifs related to homeodomain, nuclear receptor, and forkhead box family proteins (**Figure S6A**). Gene Ontology (GO) enrichment analysis revealed that BRD4 targets were strongly enriched for processes related to transcription, RNA processing, translation, and mitosis, with transcription- and mitosis-related targets exhibiting substantial repression by BET degraders (**Figure S6B,C,** and **Table S6**). PPI network construction indicated that BRD4 targets cooperatively participate in the several well-established DDLPS-relevant nodes, including ribosome biogenesis, MDM2, and chromatin assembly and regulation (**Figure S6D**)[46–51]. To visualize loop-associated regulatory mechanisms, *looplook* also provides a genome visualization component to display BRD4 binding sites, chromatin loops, and selected target gene loci (e.g., *VEGFC*, *TGFB2*, *MEIS2*, and *ARL4C*) (**Figure S6E**).

Relative to *ChIPseeker*, *looplook*-unique target genes were preferentially enriched for pathways associated with the spindle pole, DNA replication, and mitochondrial inner membrane, whereas genes commonly annotated by both tools were largely confined to fundamental cellular processes, including adhesion, translation, nuclear transport, and transcription (**Figure 4E**). Additionally, Integrative Genomics Viewer (IGV) visualization across representative oncogenic target loci supported the robust linkage of distal BRD4-bound enhancers to target TSSs (**Figure 4F,G, Figure S7**). In each case, distal enhancers marked by prominent signals of H3K27ac, H3K4me1, and ATAC-seq (but low H3K4me3) were looped to active target promoter, accompanied by strong transcriptional downregulation following ARV-825 and ZBC-260 treatment (**Figure 4F,G, Figure S7**). Taken together, these results establish *looplook* as a more robust and comprehensive framework than conventional linear annotation for cistrome analysis.

### Case study 2: *Looplook* extends the enhancer-associated FOSL2 functional network

FOSL2 (also designated Fra-2) belongs to the activator protein-1 (AP-1) family of TFs and serves as a critical promoter of DDLPS pathogenesis. As BET proteins have been shown to maintain both FOSL2 expression and FOSL2-associated transcriptional program [47, 52], we reasoned that FOSL2 targets would be vulnerable to BET protein depletion. We next tested *looplook*’s capacity to resolve the FOSL2 cistrome in LPS141 cells. Analysis of FOSL2 ChIP-seq data indicated that FOSL2 was widely recruited to non-promoter regions, with only one-third of peaks localized at promoters (**Figure 5A**), suggesting that FOSL2 is preferentially involved in distal gene regulation. Integration of 3D looping data showed that 45.6% of FOSL2 binding events (*n* = 15,818) were within loop anchors, suggesting widespread engagement in chromatin interactions (**Figure 5A**).

**Figure 5.**
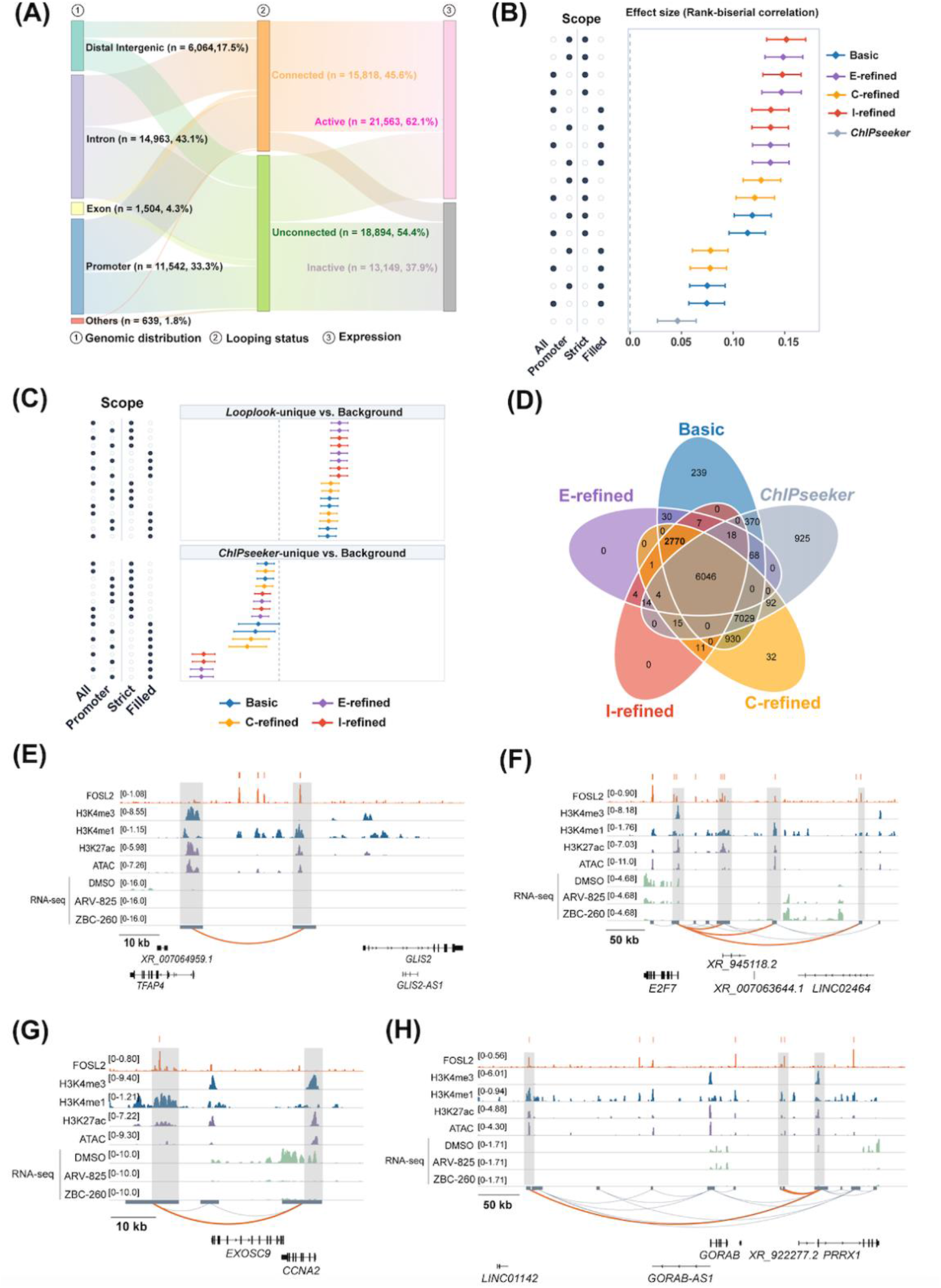
Comparative benchmarking of *looplook* against *ChIPseeker* in annotating FOSL2 targets. (A) Stacked bar plots showing the genomic distribution of FOSL2 peaks, looping status (Connected vs. Unconnected), and target gene transcriptional activity (Active vs. Inactive). (B) Effect size estimates (rank-biserial correlation) for FOSL2 target gene sets identified by *looplook* across four refinement configurations (Basic, E-refined, C-refined, and I-refined) and four peak−target annotation scopes (All: all peaks; Promoter: promoter-associated peaks; Strict: loop-engaged targets, Filled: loop-engaged targets plus fallback-filled targets), compared with the conventional linear nearest-TSS method (*ChIPseeker*) under ZBC-260 treatment. Data points represent effect size estimates with error bars indicating 95% confidence intervals. (C) Comparative effect size distributions across selected FOSL2 target gene subsets (*looplook*-unique vs. Background; *ChIPseeker*-unique vs. Background) across indicated annotation modes. (D) Venn diagram showing the overlap and unique FOSL2 target genes identified by *looplook* (Basic, E-refined, C-refined, and I-refined modes) versus *ChIPseeker*. (E−H) Linkage of distal FOSL2-bound regulatory elements to their respective target promoters in LPS141 cells.

Consistent with our analysis of the BRD4 cistrome, *looplook* outperformed *ChIPseeker* in capturing FOSL2 targets that also showed robust responsiveness to BET degraders in LPS141 cells (**Figure 5B,C, Figure S8A,B,** and **Table S5**). Moreover, permutation-based GSEA iterations conducted across multiple *looplook* operational modes yielded stable negative enrichment trajectories (**Figure S8C**−**F**), confirming that FOSL2 target genes are highly dependent on BET proteins. Under E-refined and I-refined configurations, *looplook* uniquely annotated over 2,700 FOSL2 targets, most of which engaged long-range chromatin interactions (**Figure 5D, Figure S9**). Functional annotation of *looplook*-unique FOSL2 target genes highlighted enrichment of critical pathways related to RNA splicing and ribosome, whereas genes commonly annotated by both tools were overrepresented in cell adhesion, motility, and morphogenesis processes (**Figure S10,** and **Table S6**).

Motif enrichment analysis confirmed significant overrepresentation of AP-1 consensus motifs within FOSL2-associated loop anchors (**Figure S11**). Integrative visualization of representative *looplook*-annotated FOSL2 targets (e.g., *CCNA2*, *E2F7*, *PRRX1*, and *TFAP4*) further substantiated the linkage between distal enhancers and target promoters, as well as the transcriptional sensitivity of these targets to BET degraders (**Figure 5E−H**). Collectively, these findings showcase that *looplook* provides valuable biological insight into the regulatory networks underlying TF function.

## DISCUSSION

A mechanistic understanding of gene regulation remains elusive when the genome is viewed through a purely linear lens. Deciphering target genes regulated by distal CREs is critical for translating genetic and epigenomic findings into clinically actionable insights, ranging from risk stratification to the identification of therapeutic targets. Although 3D genomics datasets have extensively delineated chromatin loop networks, these static physical conformations provide only the “topological potential” for transcriptional regulation rather than “functional certainty”. In the current study, we present *looplook*, an integrative, self-contained end-to-end R package tailored for 3D- and expression-guided CRE–target gene annotation and streamlined functional analysis. Through the case studies of the BRD4 and FOSL2 cistromes, we demonstrate *looplook*’s superior annotation performance over conventional linear annotation methods, such as *ChIPseeker*, by integrating expression data and chromatin profiles. The development of *looplook* offers a valuable new option for distilling experimental omics data into functional molecular network reconstruction.

*Looplook* applies a connected component clustering algorithm for network reconstruction, incorporating a CRE reclassification strategy (e.g., P-to-eP and G-to-eG) that preserves graph connectivity. A key direction for future development is the incorporation of loop quality metrics (such as interaction scores or read counts) as quantitative edge weights, which would further enhance the prioritization of key drivers within complex regulatory hubs compared with the current binary model. Notably, the reclassification scheme embedded in *looplook* theoretically enables unprecedented system-level detection of higher-order regulatory modules (e.g., cooperative CREs) and underpins multi-hop network diffusion (as exemplified by Cluster 2 in **Figure 2**), although this application was not explored in the current case studies. In addition, the optional multi-hop network diffusion parameter offers systems biologists a highly customizable analytical engine to quantitatively explore complex regulatory effects within clustered enhancers (e.g., super-enhancers), enhancer cliques, promoter clusters, or dense regulatory hubs[53].

Remarkably, *looplook* supports both expression- and chromatin-aware refinement for target annotation, enabling orthogonal validation of CRE reclassification. For example, by overlaying histone modification and chromatin accessibility evidence, *looplook* resolves ambiguous eP/eG classifications into more definitive anchor types (E, P, G, or Dual). The identification of dual-function CREs concurrently carrying promoter and enhancer chromatin signatures represents an important advance, as such elements have been increasingly recognized as regulatory hubs bridging promoter and enhancer functions[43, 45]. In addition, in gene-dense loci, *looplook* is likely to provide more accurate target assignment through its integrated genic biotype prioritization and dynamic topological reclassification. The current version of *looplook* implements a series of curated rules for chromatin state inference; nevertheless, we envision that further integration of machine learning-based approaches in chromatin analysis and CRE activity prediction will likely strengthen the overall accuracy and efficiency of refined target annotation.

Distinct from existing algorithms constrained by precompiled/pretrained databases or linear assignment paradigms, *looplook* supports customized analysis with user-supplied chromatin interaction, and transcriptomic and epigenomic data from any species via local, privacy-preserving computation. Furthermore, it may help to resolve long-standing challenges in the functional interpretation of non-coding variants and distal CREs by accurately prioritizing GWAS/eQTL causal target genes through chromatin loop-mediated distal regulation mapping, thereby eliminating the annotation bias inherent to linear proximity-based methods. Nevertheless, *looplook*-derived CRE–target gene linkages can be synergized with experimental validation (e.g., CRISPRi) and Mendelian randomization to strengthen causal inference.

While *looplook* is broadly applicable to various types of chromatin interactions, it may be most suitable for the study of activating CREs (particularly enhancers) and their target linkages, given that its expression-aware refinement strategy is intentionally designed to filter out silent genes. Its applicability to repressive elements (e.g., transcriptional repressors) or structural elements (e.g., insulator) remains to be established. A key future priority is the extension of *looplook* to single-cell 3D genomics and single-cell multiomics, which will be critical for dissecting regulatory dynamics in heterogeneous systems such as development, cellular plasticity, and cancer progression.

## CONCLUSION

In summary, *looplook* is a computationally efficient flexible multi-omics framework for 3D chromatin- and expression-informed CRE–target gene annotation. We anticipate that *looplook* will empower users to deploy diverse analytical workflows on custom datasets, generating robust outputs and actionable biological insights into gene regulatory networks. This framework is well suited for dissecting E–P interactions across humans, model organisms, and understudied species.

## METHODS

### Cell culture

DDLPS cell line LPS141 (RRID: CVCL_M823) was maintained in a standard culture condition[48]. For drug treatment experiments, LPS141 cells were treated with dimethyl sulfoxide (DMSO), ZBC-260 (T14550; TargetMol, Wellesley Hills, MA, USA), or ARV-825 (T5434; TargetMol).

### RNA extraction and sequencing

Total RNA was isolated from cell pellets using the RNeasy Mini Kit (Qiagen). RNA integrity was assessed prior to library preparation. Library construction and paired-end Illumina sequencing were conducted at BGI Tech Solutions Co., Ltd.

### Chromatin immunoprecipitation (ChIP) and sequencing

LPS141 cells were crosslinked with 1% formaldehyde at room temperature and subjected to nuclei extraction, sonication, and ChIP using an antibody specific to H3K27me3 (ab192985; Abcam; RRID: AB_2650559) following a standard ChIP procedure[47]. Immunoprecipitated DNA was eluted, reverse-crosslinked, purified, and used for library construction, followed by Illumina sequencing.

### Assay for transposase-accessible chromatin using sequencing (ATAC-seq)

LPS141 nuclei were isolated and resuspended in a transposition mixture containing digitonin, Tween-20, Tn5 transposase, and TD buffer from the Nextera kit (Illumina). Tagmentation was carried out for 30 min at 37°C in a ThermoMixer. The tagmented DNA was subsequently amplified, subjected to size selection, and sequenced.

### Data acquisition and preprocessing

RNA-seq data from ARV-825-treated LPS141 cells and ChIP-seq data for FOSL2, BRD4, H3K4me1, H3K4me3, and H3K27ac were obtained from GSE111254. Processed H3K27ac HiChIP data were downloaded from GSE213300 and used without re-calling. Single-end ChIP-seq reads were aligned with *Bowtie2* to hg38 reference. Alignments with MAPQ ≥ 1 were retained, and duplicates were removed using Picard *MarkDuplicates*. Fragment lengths were estimated separately for each sample using *phantompeakqualtools*. Peaks were called using *MACS2* with -f AUTO --nomodel -B --SPMR -q 0.01, with --extsize set to the sample-specific estimated fragment length. Paired-end ATAC-seq reads were aligned using *Bowtie2*, and properly paired alignments with MAPQ ≥ 30 were retained. After duplicate removal using *GATK MarkDuplicates*, peaks were called with *MACS2* using -f BAMPE --keep-dup all --call-summits -B --SPMR -q 0.01. Paired-end RNA-seq reads were aligned to the Ensembl GRCh38.106 annotation using *STAR* (v2.7.10b) with --quantMode TranscriptomeSAM GeneCounts, and quantified using RSEM. Differential expression was analyzed using *DESeq2* after retaining genes with a minimum count of 4. All analyses used the GRCh38/hg38 genome assembly and *TxDb.Hsapiens.UCSC.hg38.knownGene*. Software versions and principal parameters are provided in **Table S7**.

### *Looplook* network construction and refinement

HiChIP loops from two biological replicates were consolidated with *looplook* in consensus mode using a 500-bp anchor gap. Loops detected in both replicates were retained (*min_consensus* = 2). BRD4 or FOSL2 ChIP-seq peaks were assigned to consensus-loop anchors on GRCh38 and were required to overlap an anchor by at least 1 bp. The assignments used direct loop contacts (*neighbor_hop* = 0), the 95th percentile of unique contacts to define hubs, and biotype-first conflict resolution. The biotype priority was protein-coding, small non-coding RNA, antisense, long non-coding RNA, and pseudogene. Within the highest-priority tier, candidates with expression at least 10% of the maximum were retained. Expression refinement used DMSO-control expression, an absolute threshold of 0.5 TPM (transcripts per million), and expression-based anchor reclassification.

Chromatin refinement was performed by integrating peak tracks for H3K4me1, H3K27ac, H3K4me3, H3K27me3, and ATAC-seq, together with H3K4me1 and H3K4me3 bigWig signal profiles. Chromatin features were associated with anchors using a 200-bp gap and a minimum overlap of 100 bp. A signal ratio threshold of 3 (H3K4me1:H3K4me3) was applied, and target assignments were recomputed after refinement.

### Benchmarking against the linear nearest-TSS annotation method (*ChIPseeker*)

The benchmarking strategy involved 16 “*neighbor_hop* = 0” assignment configurations, generated by crossing four pipelines with four target scopes. The pipelines were defined as follows: Basic (initial loop-based annotation), E-refined (expression-aware refinement), C-refined (chromatin-aware refinement), and I-refined (dual refinement by expression and chromatin information). Target scopes comprised all loop-connected genes or only promoter-linked genes. Within each scope, Strict mode retained only evidence-supported loop-anchor targets, whereas Filled modes also included fallback candidates. As a comparator, *ChIPseeker* was run against *TxDb.Hsapiens.UCSC.hg38.knownGene* with a ±5-kb window around TSSs. The complete set of genes annotated by *ChIPseeker* via the nearest-gene rule was used as the reference. Genes not captured by either *looplook* or *ChIPseeker* were designated as non-assigned controls.

## AUTHOR CONTRIBUTIONS

**Ying Zhang**: Conceptualization, investigation, writing - original draft, methodology, visualization, software, formal analysis, data curation. **Xingze Huang**: Validation, methodology, software, investigation, writing - review & editing. **Huayu Chen**: Validation, investigation, writing - review & editing. **Long Xie**: Investigation, visualization, writing - review & editing. **Ye Chen**: Supervision, investigation, funding acquisition, writing - review & editing, conceptualization, resources, project administration, formal analysis, validation, data curation. **Liang Xu**: Conceptualization, investigation, funding acquisition, writing - original draft, writing - review & editing, methodology, validation, visualization, project administration, formal analysis, resources, supervision.

## ACKNOWLEDGEMENTS

We thank Prof. H. Phillip Koeffler, Prof. Jinrong Peng, Prof Shaomeng Wang, and Dr. Lingwen Ding for kind support, stimulating discussions, and constructive advice throughout the study. DeepSeek (V3) and Google Gemini were utilized as auxiliary tools to refine grammatical accuracy, enhance sentence fluency, and assist in code optimization and debugging. Graphical illustrations were created with Figdraw (www.figdraw.com). This work is funded by the National Natural Science Foundation of China (32270746 to L.X.), the National Science and Technology Major Project of the Ministry of Science and Technology of China (2025ZD0547700 to L.X.), the Zhejiang Provincial Natural Science Foundation of China (LZ24H160004 to Y.C.), and the Fundamental Research Funds for the Central Universities (226-2025-00101 to L.X.).

## ETHICS STATEMENT

No animals or humans were involved in this study.

## DATA AVAILABILITY STATEMENT

The *looplook* R package is open-source and freely available, with its source code, comprehensive documentation, and tutorials hosted on GitHub (https://github.com/zying106/looplook). The benchmarking evaluation scripts are publicly available on GitHub (https://github.com/zying106/looplook_benchmark.git). The package will also be made available through the Bioconductor repository. RNA-seq, ChIP-seq, and ATAC-seq data generated in the Xu lab have been deposited in the Genome Sequence Archive in National Genomics Data Center (https://ngdc.cncb.ac.cn/gsa) under HRA020868. The related processed files have been deposited in the OMIX database (https://ngdc.cncb.ac.cn/omix: accession no. OMIX020322).

