## Supporting Information for "*Looplook*: Integrating multiomics refinement and graph clustering for target assignment and functional inference of chromatin regulatory networks"

**This file includes:**

**Extended Methods**

**Figures S1 to S11**

**Tables S1 to S3**

### Extended Methods

#### Cell culture

Human DDLPS cell line LPS141 (RRID: CVCL_M823) was maintained in Roswell Park Memorial Institute 1640 medium supplemented with 10% fetal bovine serum (F101-01; Vazyme, Nanjing, China) and 1% penicillin-streptomycin (ST488S; Beyotime, Shanghai, China). Cells were cultured at 37°C in a humidified incubator containing 5% $CO_2$. For drug treatment experiments, LPS141 cells were treated with dimethyl sulfoxide (DMSO), ZBC-260 (T14550; TargetMol, Wellesley Hills, MA, USA), or ARV-825 (T5434; TargetMol).

#### Western blotting

Cells were harvested and incubated for 20 min on ice in whole-cell lysis buffer containing 50 mM Tris-HCl (pH 8.0), 420 mM NaCl, 5% glycerol, 0.1% NP-40, and 0.1 mM EDTA. Immediately before use, the buffer was supplemented with 1 mM dithiothreitol, 1 mM phenylmethylsulfonyl fluoride, 1× protease inhibitor cocktail (C0001; TargetMol), 1× phosphatase inhibitor cocktail (C0004; TargetMol), 1 mM $MgCl_2$, and BenzoNuclease (1:500; M056; Novoprotein, Suzhou, China). Following protein quantification using the Bradford assay, equal amounts of protein were resolved by sodium dodecyl sulfate–polyacrylamide gel electrophoresis and analyzed by immunoblotting according to standard procedures. Primary antibodies against BRD2 (5848; Cell Signaling Technology; RRID: AB_10835146), BRD3 (11859-1-AP; Proteintech; RRID: AB_2065902), BRD4 (A700-004; Bethyl; RRID: AB_2631885), FOSL2 (19967; Cell Signaling Technology; RRID: AB_2722526), and β-ACTIN (66009-1-Ig; Proteintech; RRID: AB_2687938) were used. β-ACTIN served as the loading control.

#### Chromatin immunoprecipitation (ChIP) and sequencing

LPS141 cells were crosslinked with 1% formaldehyde at room temperature and then washed three times with ice-cold phosphate-buffered saline. Nuclei were isolated, resuspended in SDS lysis buffer, and incubated on ice for 10 min. Chromatin was sheared with a Bioruptor sonicator (Diagenode) to obtain DNA fragments of approximately 200–500 bp. The soluble chromatin fraction was incubated overnight with Dynabeads Protein A/G pre-conjugated to H3K27me3-specific antibodies (ab192985; Abcam; RRID: AB_2650559). The magnetic beads were washed sequentially with ice-cold low-salt wash buffer, high-salt wash buffer, LiCl wash buffer, and TE buffer. Immunoprecipitated DNA was eluted, reverse-crosslinked, and purified using a QIAquick PCR Purification Kit (Qiagen). Library construction was performed using the ThruPLEX DNA-seq Kit (R400406; Takara), and subjected to Illumina sequencing.

**Assay for transposase-accessible chromatin using sequencing (ATAC-seq)**

For each assay, 50,000 viable LPS141 cells were resuspended in 50 μL of resuspension buffer containing 10 mM Tris-HCl (pH 7.4), 10 mM NaCl, 3 mM $MgCl_2$, 0.1% NP-40, 0.1% Tween-20, 0.01% digitonin, and protease inhibitors. Nuclei were isolated and resuspended in a transposition mixture containing digitonin, Tween-20, Tn5 transposase, and TD buffer from the Nextera kit (Illumina). Tagmentation was performed for 30 min at 37°C with mixing at 1,000 rpm in a ThermoMixer. The reaction was stopped with PB buffer, and DNA was purified using a MinElute Kit (Qiagen). Tagmented DNA was then amplified, size-selected, and sequenced.

**Functional enrichment analyses and statistics**

Differentially expressed genes were ranked by $\log_2$(fold change). For effect-size analyses, the response score was defined as $-\log_2$(fold change), such that positive values indicated stronger downregulation after treatment. Group differences were summarized using rank-biserial correlation derived from the Wilcoxon rank-sum statistic. Ninety-five percent confidence intervals were estimated from 2,000 stratified bootstrap replicates using bias-corrected and accelerated intervals.

GSEA was performed with *clusterProfiler::GSEA* using the following parameters: *pvalueCutoff* = 1.1, *pAdjustMethod* = none, *minGSSize* = 10, *maxGSSize* = 50,000, and deterministic seeding. Each mode was evaluated in 300 resampling iterations with a requested sample size of 200 genes per group. When a pool contained fewer than 200 genes, the script sampled 80% of the available genes, retained at least 10 genes, and matched group sizes to the smaller eligible pool. The unique-set analysis separately sampled *looplook*-only, shared, and *ChIPseeker*-only genes at a common size of up to 200. Normalized enrichment scores (NES) were reported as density distributions and cumulative moving averages.

GO enrichment was performed with *clusterProfiler::enrichGO* and *org.Hs.eg.db* with the biological process ontology. Enriched terms were retained at a significance threshold of $P < 0.01$; all ranked genes successfully mapped to Entrez identifiers served as the universe. Motif scanning was performed against the JASPAR2020 CORE collection with *TFBSTools* and *motifmatchr* with a threshold of $1\times10^{-4}$. Motif enrichment was tested against GC-matched background regions by one-sided Fisher's exact tests, and results were reported as Benjamini–Hochberg false discovery rate (FDR) and $\log_2$(odds ratio).

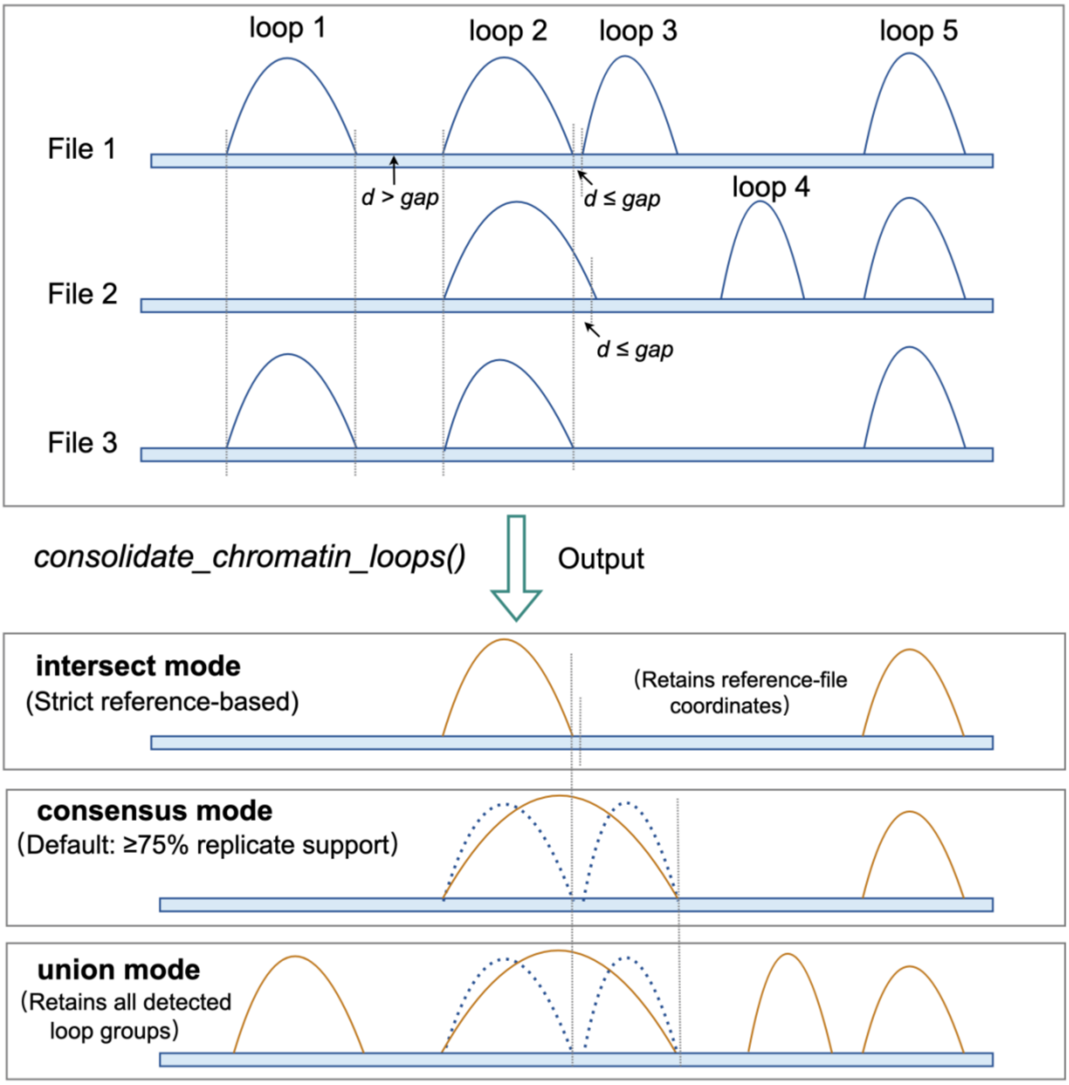


**Figure S1. Schematic of the topology-merging engine in the Multi-Source Consensus module**. The *consolidate_chromatin_loops* function identifies consistent topological features across heterogeneous input files by comparing loop distances ($d$) against a user-defined gap threshold. Adjacent loops with $d$ > gap are considered independent, whereas those with $d$ ≤ gap are interpreted as putatively identical biological interactions exhibiting technical variance (wobble) and are thus clustered together. This engine supports three flexible merging strategies: (1) intersect mode, which maximizes specificity by retaining only loops with fully overlapping anchors; (2) consensus mode (default), which outputs loops supported by the majority of replicates (> 75% by default); and (3) union mode, which retains all detected loops to maximize topological coverage.

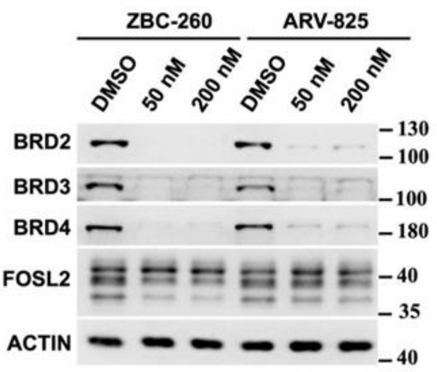


**Figure S2. Effect of ZBC-260 and ARV-825 on target proteins in LPS141 cells.** LPS141 cells were treated with indicated compounds for 8 h prior to immunoblotting analysis.

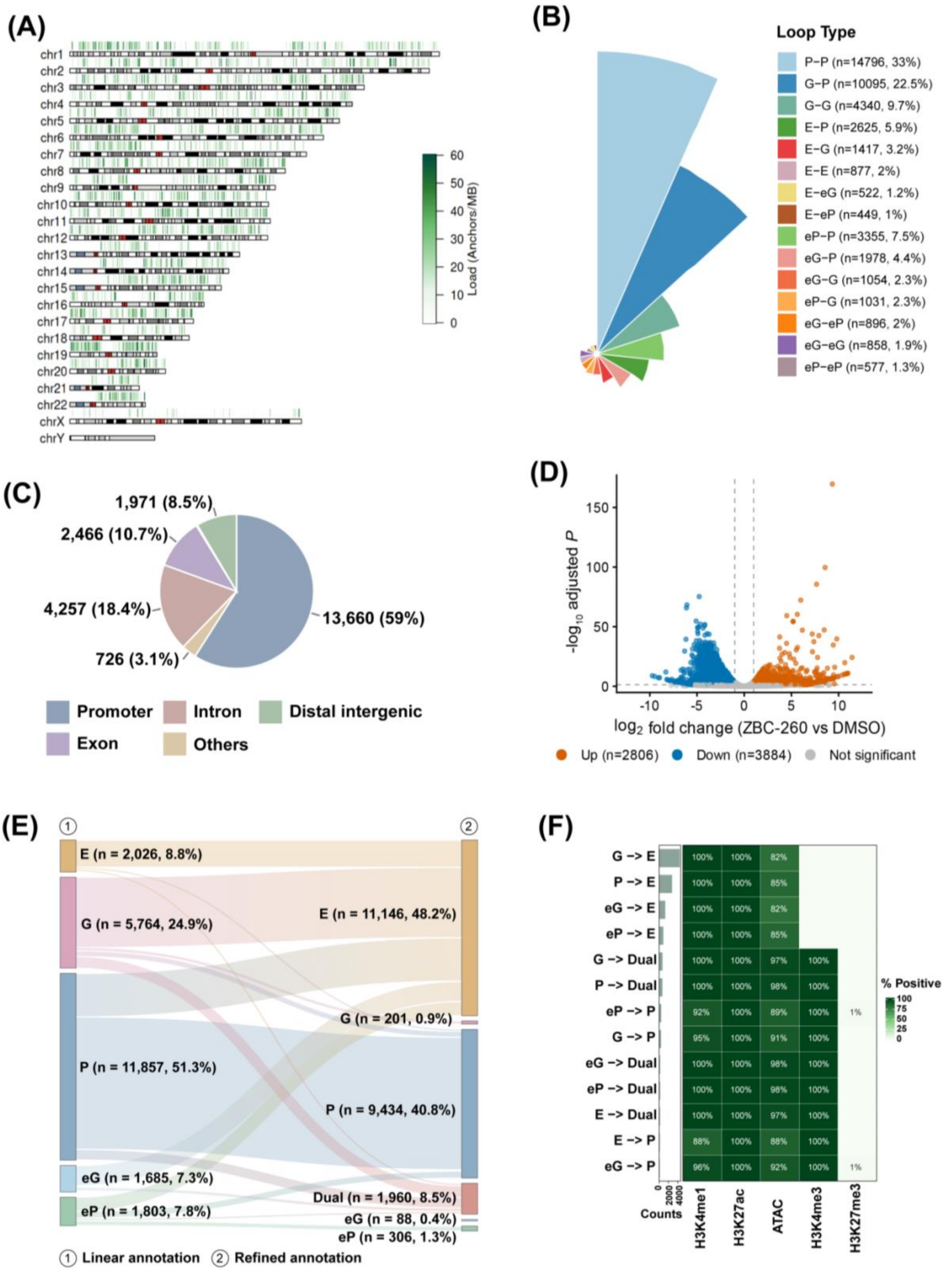


**Figure S3. Integrative chromatin and transcriptomic analysis in LPS141 cells.** (A) Chromosome ideogram showing the genome-wide density and distribution of H3K27ac HiChIP loop anchors across all chromosomes, confirming uniform coverage with a median load of ~20 anchors/MB. (B) Types of H3K27ac HiChIP loop-associated interactions. (C) Genome-wide distribution of anchors from the consolidated H3K27ac HiChIP loop set in

LPS141 cells. Pie chart showing the genomic annotation of anchor regions from the consolidated loop set across promoters, introns, distal intergenic regions, exons, and other genomic regions, with anchor counts and percentages indicated. (D) Volcano plot showing the impact of ZBC-260 treatment (200 nM, 8 h) on transcriptome in LPS141 cells. (E) Sankey diagram illustrating the CRE reclassification based on 3D looping-assisted refinement with *looplook*. Left: linear annotation of CRE; right: refined annotation. E, enhancer; P, promoter, G, gene body; eG, enhancer-like CRE in gene body; eP, enhancer-like CRE in promoter; Dual, CRE with dual functions as promoter and enhancer. (F) Heatmap showing the positive percentage of histone modifications (H3K4me1, H3K27ac, H3K4me3, and H3K27me3) and chromatin accessibility (ATAC) signals across the reclassified anchor categories. Anchor counts are shown as well.

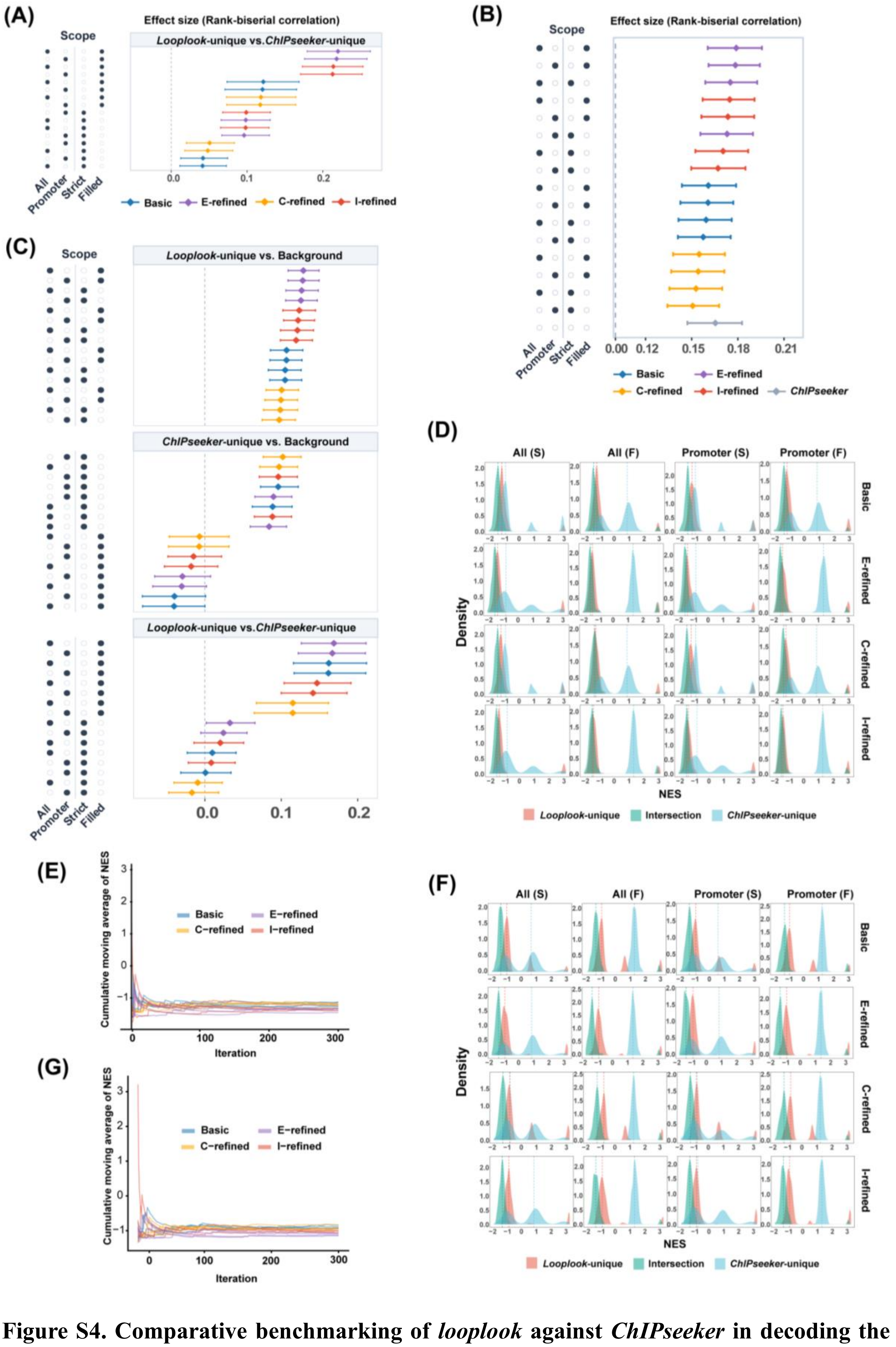


**Figure S4. Comparative benchmarking of *looplook* against *ChIPseeker* in decoding the BRD4 cistrome.** (A) Comparative effect size distributions across selected BRD4 target gene subsets (*looplook*-unique vs. *ChIPseeker*-unique) across indicated annotation modes. RNA-seq data were obtained from LPS141 cells following ZBC-260 treatment (200 nM, 8 h). (B) Effect size estimates (rank-biserial correlation) for BRD4 target gene sets identified by *looplook* across four refinement configurations (Basic, E-refined, C-refined, and I-refined) and four peak–target annotation scopes (All: all peaks; Promoter: promoter-associated peaks; Strict: loop-engaged targets, Filled: loop-engaged targets plus fallback-filled targets), compared with *ChIPseeker* under ARV-825 treatment (200 nM, 24 h). Data points represent effect size estimates with error bars indicating 95% confidence intervals. (C) Comparative effect size distributions across selected BRD4 target gene subsets (*looplook*-unique vs. Background; *ChIPseeker*-unique vs. Background; *looplook*-unique vs. *ChIPseeker*-unique) across indicated annotation modes. RNA-seq data were obtained from LPS141 cells following ARV-825 treatment (200 nM, 24 h). (D) Density distributions of Normalized Enrichment Scores (NES) across multiple execution modes and fallback strategies comparing *looplook*-unique, intersection, and *ChIPseeker*-unique BRD4 target gene sets under ZBC-260 treatment. (E) Cumulative moving average of NES across permutation iterations under ZBC-260 treatment, showing rapid convergence and sustained negative enrichment for *looplook*-unique BRD4 target gene sets. (F) Density distributions of NES across multiple execution modes and fallback strategies comparing *looplook*-unique, intersection, and *ChIPseeker*-unique BRD4 target gene sets under ARV-825 treatment. (G) Cumulative moving average of NES across permutation iterations under ARV-825 treatment.

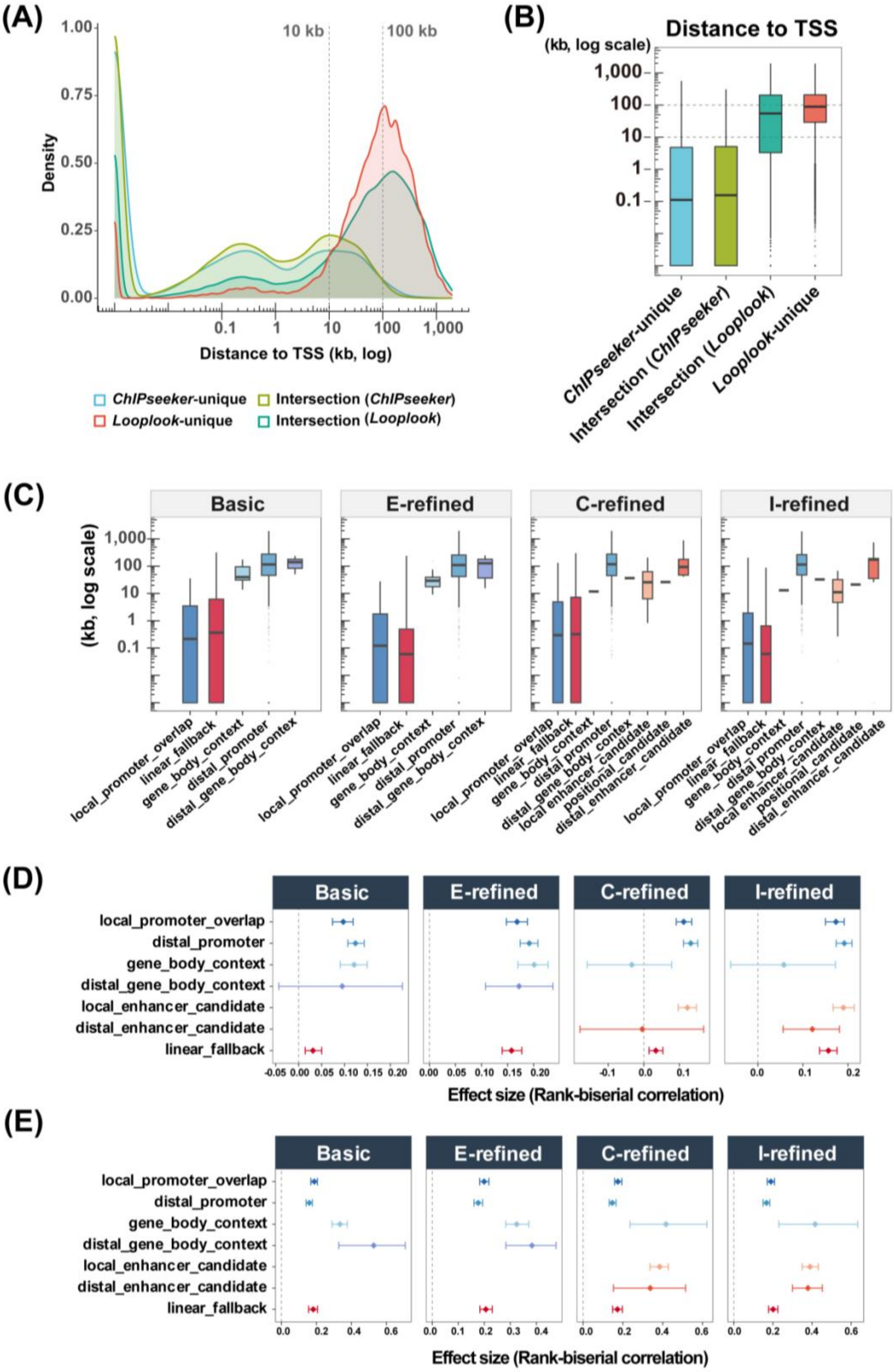


**Figure S5. *Looplook* is advantageous in establishing distal CRE−BRD4 target gene linkages.** (A) Density plot showing the genomic distance between CREs and cognate target TSSs (Y-axis in log scale) for linkages from *ChIPseeker*-unique, *looplook*-unique, and intersection BRD4 target gene sets. (B) Boxplots showing the genomic distance between CREs and cognate target TSSs (Y-axis in log scale) for linkages from *ChIPseeker*-unique, intersection, and *looplook*-unique BRD4 target gene sets. (C) Genomic distance between CREs and cognate target TSSs across sub-categories of BRD4 target gene sets, including those involving *local_promoter_overlap* (CRE−local promoter overlap), *distal_promoter* (CRE–distal promoter link), *distal_gene_body_context* (CRE–distal gene body link), *gene_body_context* (CRE−gene body overlap), *local_enhancer_candidate* (CRE–local enhancer candidate), *distal_enhancer_candidate* (CRE–distal enhancer candidate), *positional_candidate* (CRE–position-based candidate), *linear_fallback* (CRE linear fallback),. Results annotated by *looplook* under Basic, E-refined, C-refined, and I-refined configurations are shown. (D,E) Effect size evaluations under (D) ZBC-260 or (E) ARV-825 treatment across sub-categories of BRD4 target gene sets as described in (C).

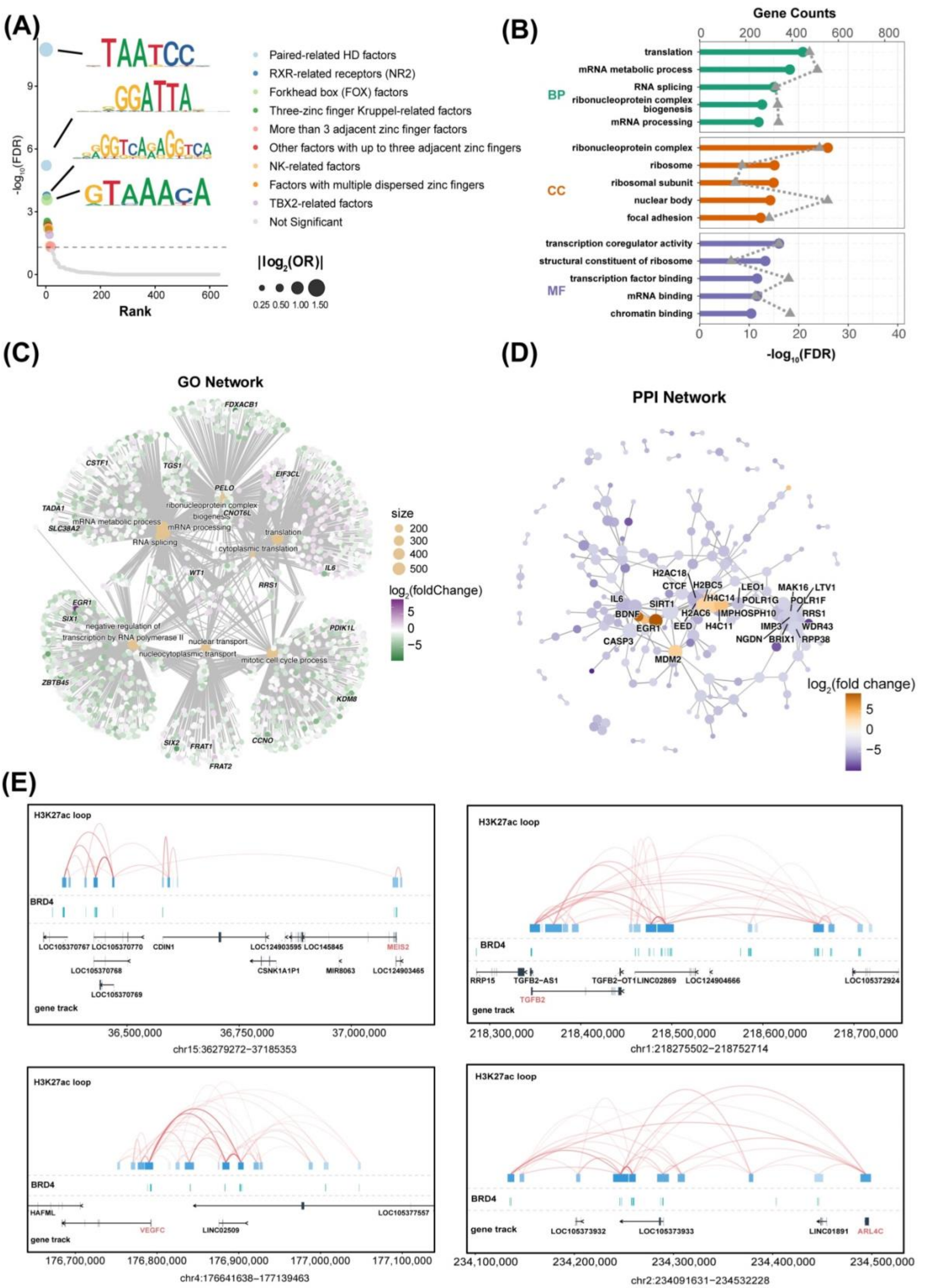


**Figure S6. Application of streamlined bioinformatics tools embedded in *looplook*.** (A) Rank-ordered transcription factor motif enrichment analysis for *looplook*-resolved BRD4 binding sites. Top enriched motifs and their corresponding TF families are highlighted. OR, odds ratio. (B,C) GO enrichment analysis of BRD4 targets annotated by *looplook*. (D) PPI network construction of BRD4 targets annotated by *looplook* (*ppi_score* = 500). (E) Integrative visualization of BRD4 binding sites and chromatin loops associated with selected BRD4 target gene loci.

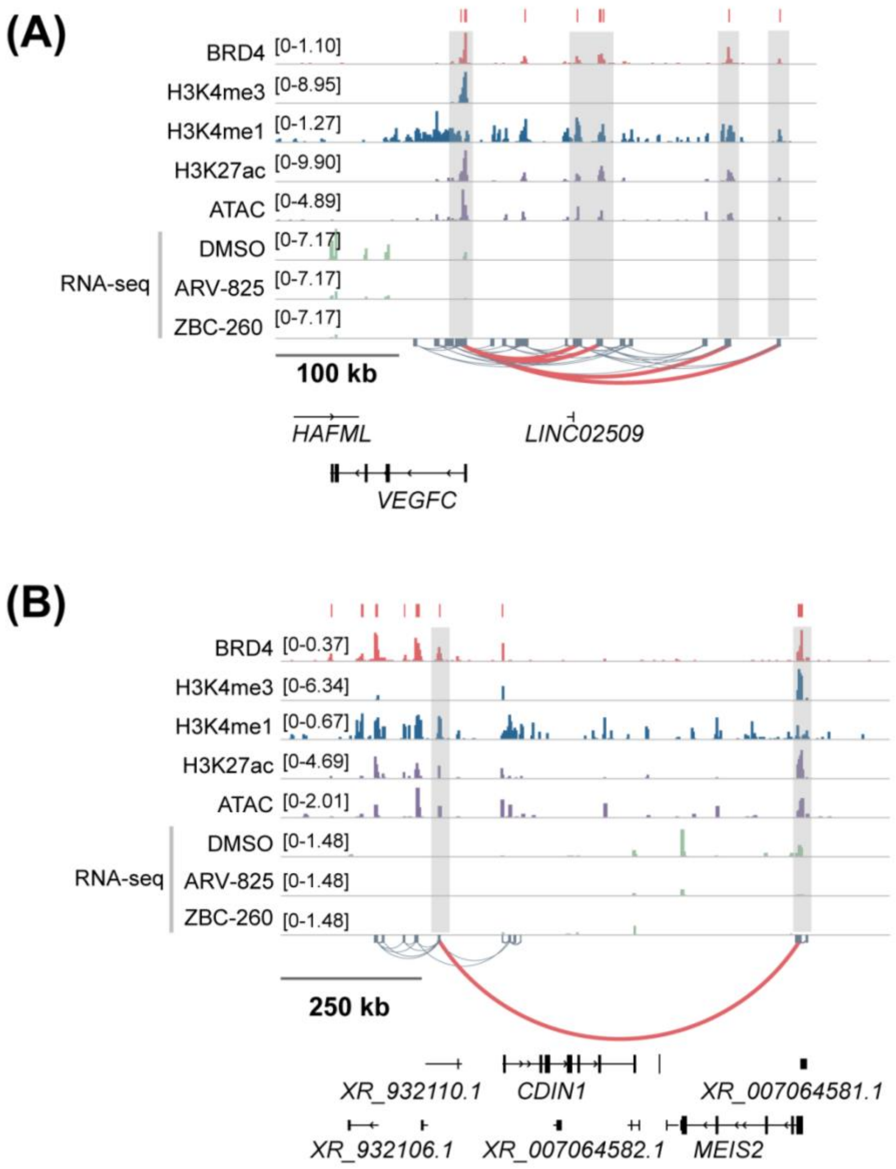


**Figure S7. Linkage of distal BRD4-bound regulatory elements to the *VEGFC* and *MEIS2* loci in LPS141 cells.**

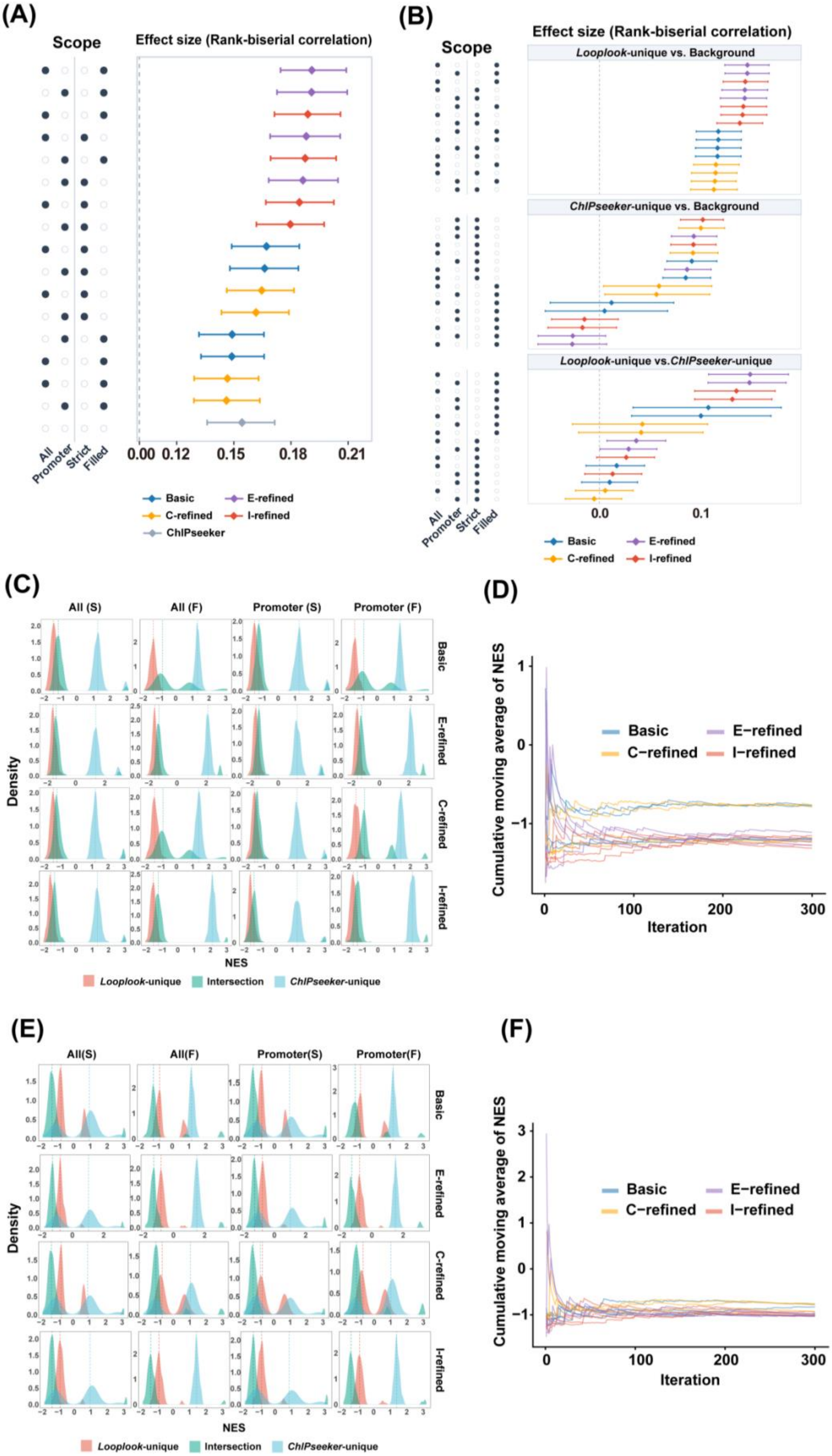


**Figure S8. Comparative analysis of *looplook* versus *ChIPseeker* in decoding the FOSL2**

**cistrome.** (A) Effect size estimates (rank-biserial correlation) for FOSL2 target gene sets identified by *looplook* across four refinement configurations (Basic, E-refined, C-refined, and I-refined) and four peak−target annotation scopes (All: all peaks; Promoter: promoter-associated peaks; Strict: loop-engaged targets, Filled: loop-engaged targets plus fallback-filled targets), compared with *ChIPseeker* under ARV-825 treatment (200 nM, 24 h). Data points represent effect size estimates with error bars indicating 95% confidence intervals. (B) Comparative effect size distributions across selected FOSL2 target gene subsets (*looplook*-unique vs. Background; *ChIPseeker*-unique vs. Background; *looplook*-unique vs. *ChIPseeker*-unique) across indicated annotation modes. RNA-seq data were obtained from LPS141 cells following ARV-825 treatment (200 nM, 24 h). (C) Density distributions of NES across multiple execution modes and fallback strategies comparing *looplook*-unique, intersection, and *ChIPseeker*-unique FOSL2 target gene sets under ZBC-260 treatment. (D) Cumulative moving average of NES across permutation iterations under ZBC-260 treatment, showing rapid convergence and sustained negative enrichment for *looplook*-unique FOSL2 target gene sets. (E) Density distributions of NES across multiple execution modes and fallback strategies comparing *looplook*-unique, intersection, and *ChIPseeker*-unique FOSL2 target gene sets under ARV-825 treatment. (F) Cumulative moving average of NES across permutation iterations of FOSL2 target gene sets under ARV-825 treatment.

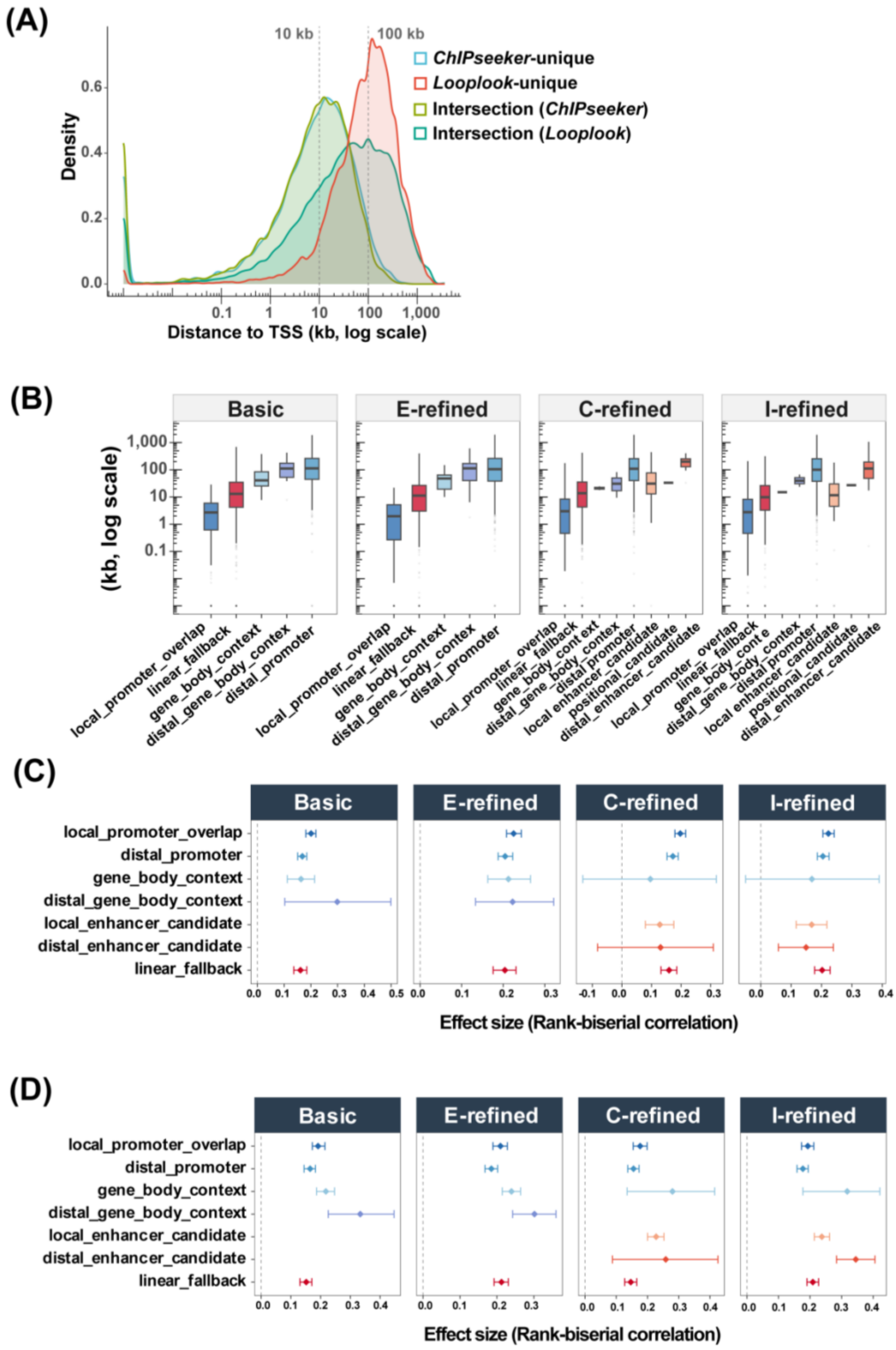


**Figure S9. *Looplook* is advantageous in establishing distal CRE−FOSL2 target gene linkages.** (A) Density plot showing the genomic distance between CREs and cognate target TSSs (Y-axis in log scale) for linkages from *ChIPseeker*-unique, *looplook*-unique, and intersection FOSL2 target gene sets. (B) Genomic distance between CREs and cognate target TSSs across sub-categories of FOSL2 target gene sets, including *local_promoter_overlap* (CRE−local promoter overlap), *distal_promoter* (CRE–distal promoter link), *distal_gene_body_context* (CRE–distal gene body link), *gene_body_context* (CRE−gene body overlap), *local_enhancer_candidate* (CRE–local enhancer candidate), *distal_enhancer_candidate* (CRE–distal enhancer candidate), *positional_candidate* (CRE–position-based candidate), *linear_fallback* (CRE linear fallback),. Results annotated by *looplook* under Basic, E-refined, C-refined, and I-refined configurations are shown. (C,D) Effect size evaluations under (C) ZBC-260 or (D) ARV-825 treatment across sub-categories of FOSL2 target gene sets as described in (B).

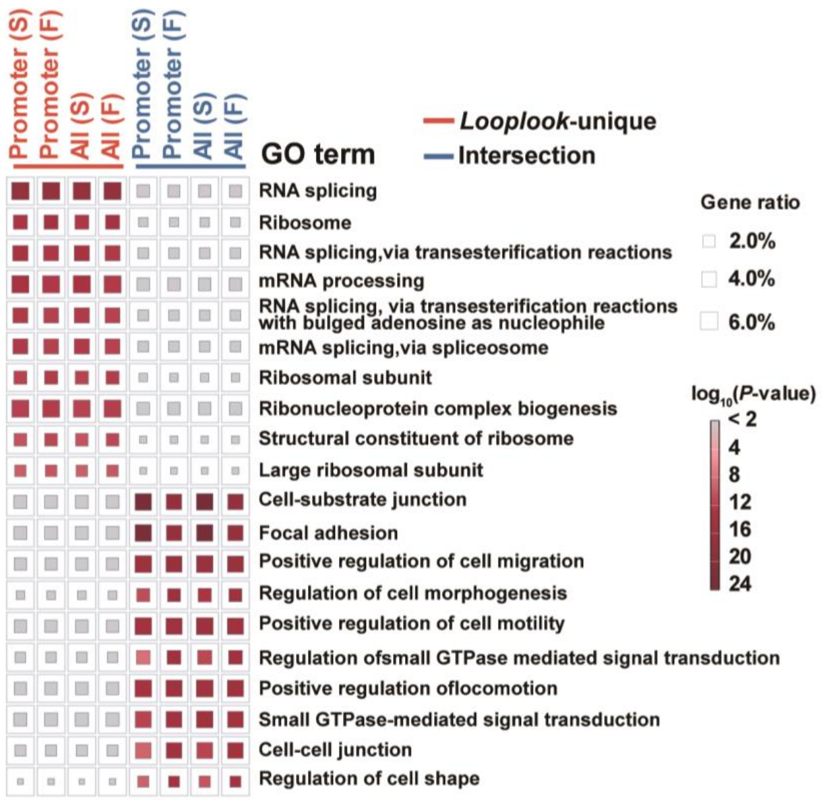


**Figure S10. GO enrichment analysis comparing *looplook*-unique and commonly annotated (intersection) FOSL2 target genes.** Dot sizes represent gene ratios; colors indicate statistical significance ($-\log_{10} P$-value) across multiple peak−target scopes (S, strict; F, fallback-filled).

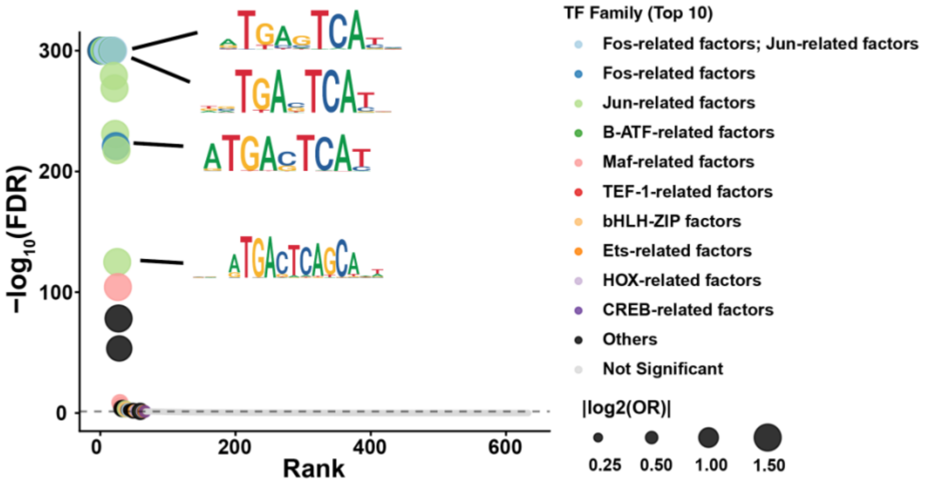


**Figure S11. Rank-ordered transcription factor motif enrichment analysis for *looplook*-resolved FOSL2 peaks.**

**Table S1. Chromatin state inference hierarchy (first-match priority).**

| Priority | Chromatin state | Mark combination | Biological interpretation |
|---|---|---|---|
| 1 | *conflicting_marks* | H3K27me (+) & any active mark | Bivalent or poised chromatin |
| 2 | *dual_like* | H3K4me1 (+) & H3K4me3 (+) | Dual function as promoter & enhancer |
| 3 | *active_enhancer_like* | H3K4me1 (+) & H3K27ac (+) & ATAC (+) & H3K4me3 (–) | Active enhancer |
| 4 | *intermediate_enhancer_like* | H3K4me1 (+) & H3K4me3 (–) & (H3K27ac (+) or ATAC (+)) | Primed enhancer with partial activation evidence |
| 5 | *other_enhancer_like* | H3K4me1 (+) or H3K27ac (+) | Weak or context-dependent enhancer |
| 6 | *promoter_like* | H3K4me3 (+) | Active or poised promoter |
| 7 | *repressed* | H3K27me3 (+) only | Inactive region |
| 8 | *uncertain* | No definitive signature present | Uncertain activity |
| 9 | *no_data* | No information available | |

NOTE: Each mark at an anchor is recorded as '+' (at least one peak of that mark overlaps the anchor) or '–' (synonymous with *not_called*, indicating the mark was analyzed but no peak was detected at the anchor). Only '–' is treated as negative evidence. These *chromatin_state* labels are descriptive; the final anchor type is determined by the reclassification rules, and *dual_like* anchors are resolved into the final anchor type based on the H3K4me1/H3K4me3 bigWig signal ratio, as specified in the reclassification rules.

**Table S2. Anchor-type reclassification rules based on chromatin status.**

| Tier | Positional type | Chromatin status | Outcome |
|---|---|---|---|
| A. Curated override | any | within *enhancer_bed*, H3K4me3 (+) | Dual |
| | any | within *enhancer_bed*, H3K4me3 (–) | E |
| B. Silent (eP/eG) | eP/eG | dual-like, H3K4me1/H3K4me3 signal ratio | Dual/keep/P |
| | eP/eG | H3K4me3 (+) & H3K4me1 (–) | P |
| | eP/eG | *canonical/strong* enhancer evidence and *active/intermediate enhancer* | E |
| | eP/eG | *promoter_like* | P |
| | any | *conflicting_marks* (bivalent/poised) | unchanged |
| C. Expressed (P/E/G) | P/E/G | dual-like, H3K4me1/H3K4me3 signal ratio | Dual/keep/P |
| | P | H3K4me1 (+) & H3K4me3 (–) & (H3K27ac (+) or ATAC (+)) | E |
| | E/G | H3K4me3 (+) & H3K4me1 (–) | P |
| | G | H3K4me1 (+) & H3K4me3 (–) & (H3K27ac (+) or ATAC (+)) | E |
| D. Fallback | any | atypical | unchanged |

NOTE: Rules are tiered by data priority: A (curated *enhancer_bed* file), B (positional type eP/eG), C (positional type P/E/G), D (Fallback). +, detected; –, not detected. Minimal input: H3K4me1 and H3K4me3 signals; optional data: H3K27ac, ATAC, and H3K27me3. Anchors dually positive for H3K4me1 and H3K4me3 are resolved by H3K4me1/H3K4me3 bigWig ratio vs. the configurable *bw_ratio_threshold* (default 3): enhancer-dominant when ratio ≥ threshold → Dual; unresolved when no bigWig available → keep; promoter-dominant when ratio < threshold → P. The curated *enhancer_bed* overrides all signal-based rules. Anchors with *conflicting_marks* (bivalent/poised) are retained unchanged. The *enhancer_evidence* metric is a five-level confidence factor derived from the mark set by stricter criteria than the state labels: canonical (see Supplementary Table S3 for details).

**Table S3. Comprehensive anchor-type reclassification rules.**

| Priority | Positional type | Chromatin condition | Reclassified type | Rationale / note |
|---|---|---|---|---|
| 1 | any | anchor within *enhancer_bed* & H3K4me3 (+) | Dual | curated enhancers from databases; override all signal-based rules |
| 2† | any | anchor within *enhancer_bed* & H3K4me3 (–) | E | curated enhancers, without H3K4me3 signals |
| 3 | eP/eG | *dual_like* & ratio = *me1_dominant* | Dual | bigWig H3K4me1/H3K4me3 ≥ 3 → dual function as E & P |
| 4 | eP/eG | *dual_like* & ratio = unresolved | eP/eG | no bigWig available → keep |
| 5 | eP/eG | *dual_like* & ratio = *not_me1_dominant* | P | H3K4me3 dominates → P |
| 6 | eP/eG | H3K4me3 (+) & (H3K4me1 (–)) | P | H3K4me3 without H3K4me1 |
| 7 | eP/eG | enhancer_evidence ∈ {*canonical*,*strong*} & chromatin_state ∈ {*active_enhancer_like*, *intermediate_enhancer_like*} | E | orthogonal active enhancer confirmation |
| 8 | eP | chromatin_state = *promoter_like* | P | generic H3K4me3-only P |
| 9 | eG | chromatin_state = *promoter_like* | P | generic H3K4me3-only P |
| 10 | any | chromatin_state = *conflicting_marks* | unchanged | Bivalent/poised → keep; no confident call |
| 11 | P | H3K4me1 (+) & H3K4me3 (+) & ratio = *me1_dominant* | Dual | promoter-localized CREs with dual function as P & E |
| 12 | P | H3K4me1 (+) & H3K4me3 (+) & ratio = *unresolved* | P | no bigWig available → keep |
| 13 | P | H3K4me1 (+) & H3K4me3 (+) & ratio = *not_me1_dominant* | P | keep as P |
| 14† | P | H3K4me1 (+) & H3K4me3 (–) & (H3K27ac (+) or ATAC (+)) | E | lost H3K4me3, gained enhancer marks → distal E |
| 15 | E | H3K4me1 (+) & H3K4me3 (+) & ratio = *me1_dominant* | Dual | distal CREs with dual function as P & E |
| 16 | E | H3K4me1 (+) & H3K4me3 (+) & ratio = *unresolved* | E | no bigWig available → keep |
| 17 | E | H3K4me1 (+) & H3K4me3 (+) & ratio = *not_me1_dominant* | P | unannotated P |
| 18 | E | H3K4me3 (+) & H3K4me1 (–) | P | unannotated P |
| 19 | G | H3K4me1 (+) & H3K4me3 (+) & ratio = *me1_dominant* | Dual | gene-body CREs with dual function as P & E |
| 20 | G | H3K4me1 (+) & H3K4me3 (+) & ratio = *unresolved* | G | no bigWig available → keep |
| 21 | G | H3K4me1 (+) & H3K4me3 (+) & ratio = *not_me1_dominant* | P | internal promoter |
| 22 | G | H3K4me3 (+) & H3K4me1 (–) | P | internal promoter within gene body |
| 23† | G | H3K4me1 (+) & H3K4me3 (–) & (H3K27ac (+) or ATAC (+)) | E | conserved intronic enhancer |
| 24 | any | TRUE (Fallback) | unchanged | no reclassification |

† H3K4me3 is used as negative evidence.
